# Concert: Genome-wide prediction of sequence elements that modulate DNA replication timing

**DOI:** 10.1101/2022.04.21.488684

**Authors:** Yang Yang, Yuchuan Wang, Yang Zhang, Jian Ma

## Abstract

Proper control of replication timing (RT) is of vital importance to maintain genome and epigenome integrity. However, the genome-wide sequence determinants regulating RT remain unclear. Here, we develop a new machine learning method, named Concert, to simultaneously predict RT from sequence features and identify RT-modulating sequence elements in a genome-wide manner. Concert integrates two functionally cooperative modules, a selector, which performs importance estimation-based sampling to detect predictive sequence elements, and a predictor, which incorporates bidirectional recurrent neural networks and self-attention mechanism to achieve selective learning of longrange spatial dependencies across genomic loci. We apply Concert to predict RT in mouse embryonic stem cells and multiple human cell types with high accuracy. The identified RT-modulating sequence elements show novel connections with genomic and epigenomic features such as 3D chromatin interactions. In particular, Concert reveals a class of RT-modulating elements that are not transcriptional regulatory elements but are enriched with specific repetitive sequences. As a generic interpretable machine learning framework for predicting large-scale functional genomic profiles based on sequence features, Concert provides new insights into the potential sequence determinants of RT.

## Introduction

As a fundamental genome function of eukaryotic cells, the replication timing (RT) program has a highly regulated temporal pattern along the genome, which is intertwined with genome structure and other genome functions [1–3]. High-throughput Repli-seq data together with the 3D genome features derived from Hi-C have shown that the active domains towards the nuclear interior generally have early RT patterns, which are also evolutionarily conserved [3–5]. Methods have also been developed to predict RT in human cells based on epigenomic features [6, 7] and yet genomic sequence features that may modulate RT patterns remain mostly unclear [3]. Sima et al. [8] recently made a significant step forward in identifying specific *cis*-acting elements regulating early RT in mouse embryonic stem cells (mESCs). Based on CRISPR-mediated deletions on a few genomic loci, early replicating control elements (ERCEs) were experimentally validated to regulate early RT in mESCs [8]. However, it is unclear whether there are sequence features that are predictive of the entire RT program genome-wide across different cell types. We currently lack effective computational models that can accurately predict detailed genome-wide RT profiles from sequences.

In recent years, there have been a number of machine learning methods that use DNA sequences to predict functional genomic signatures as well as chromatin folding patterns. Starting with early successes of applying deep neural network based architectures to regulatory genomics [9, 10], models have been developed to generate features from the sequences to predict transcription factor (TF) binding and other short-range, local epigenetic features based on both convolution neural networks (CNNs) and recurrent neural networks (RNNs) [11–17]. More recently, a few models have been proposed to use dilated convolutions to learn long-range spatial dependencies in DNA sequences, which are critical for the prediction of long-range gene regulation and chromatin interaction [18–21]. For example, Basenji [18] uses CNNs and dilated convolutions to predict gene expressions from DNA sequences at 128bp resolution. The very recently developed Enformer increases the receptive field width as compared to Basenji and predicts gene expressions from DNA sequences by utilizing self-attention mechanism to capture long-range interaction information [22]. Enformer also uses CNNs for sequence feature extraction at the resolution of 128bp. However, for data types with lower resolution and with larger-scale signal profiles (e.g., the RT data with 5-50Kb bin size), it is particularly challenging to apply CNNs for efficient high-quality feature extraction from each genomic bin. Importantly, for long-range spatial dependency, the receptive field width of dilated convolutions is still limited in size compared to the input sequence length. In particular, the dilated convolutions use regular gaps to capture spatial dependency, which may lose information on more complicated dependency patterns (e.g., combinatorial patterns of cooperative elements at different loci). Therefore, it remains an unsolved problem to capture dependency among arbitrary loci over a very large-scale genomic region such as replication domains, which are typically larger than 500Kb in size [4], and achieve high model interpretability.

Here, we develop an interpretable, context-of-sequences-aware model for RT prediction, named Concert(CONtext-of-sequenCEs for Replication Timing), using DNA sequence features only. Concert unifies (i) modeling of long-range spatial dependencies across different genomic loci and (ii) detection of a subset of genomic loci that are predictive of the target genomic signals over large-scale spatial domains. Applications of Concert to 10 human cell lines and mESCs demonstrate that the model achieves high RT prediction accuracy. Crucially, our method can identify sequences that are potentially important for shaping the RT program (i.e., RT-modulating elements). Concert provides a generic interpretable framework for predicting potential sequence determinants for large-scale genomic signals.

## Results

### Overview of the Concert model

Concert is an interpretable, context-attentive model that is able to simultaneously identify predictive sequence elements for modulating RT and predict genome-wide RT profiles using DNA sequence features only. Specifically, Concert is structured with two functionally cooperative modules: (i) selector and (ii) predictor, which are trained jointly within one unified framework in modeling long-range spatial dependencies across genomic loci, detecting predictive sequence elements, and learning context-aware sequence feature representations (see **Methods** and **Supplementary Methods** A.1). The selector aims to estimate which genomic loci are of potential importance in modulating the target signals (RT signals in this work), approximately selecting a set of predictive sequence elements by importance estimation-based subset sampling. Leveraging sequence importance estimation from the selector, the predictor then performs selective learning of spatial dependencies across the genomic loci and makes prediction of target signals over large-scale spatial domains.

An overview of the Concert model is shown in **Fig**. 1a. The input to the model contains DNA sequences of the genomic loci within each context window. The output includes both the predicted signals and the locus-wise estimated importance scores. The estimated scores can be further processed to delineate predictive sequence elements that are important for modulating the target signals. A notable advance of Concert is that it embeds the interpretation mechanism into predictive model training, thereby not requiring separate post-hoc steps for model explanation. Importantly, the model is also capable of handling different sizes of context without increasing the model parameters, making it more scalable to the context size of genomic loci. Details of the model architecture are described in **Methods**. We applied Concert to predict RT and identify RT-modulating elements in mESCs and 10 human cell lines, including H1-hESC, H9-hESC, GM12878, HCT116, HEK293, K562, IMR-90, RPE-hTERT, U2OS, and WTC11. We used the two-fraction E/L Repli-seq data (see **Methods** for details on data processing). We only included autosomal chromosomes in the analysis, in consideration of the sex identity differences of the cell lines and the potential bias induced by the sex chromosomes. We divided the genome into consecutive non-overlapping 5Kb bins (genomic loci). Note that the processed RT signal at each locus is a continuous value, with higher value representing earlier replication.

**Figure 1:**
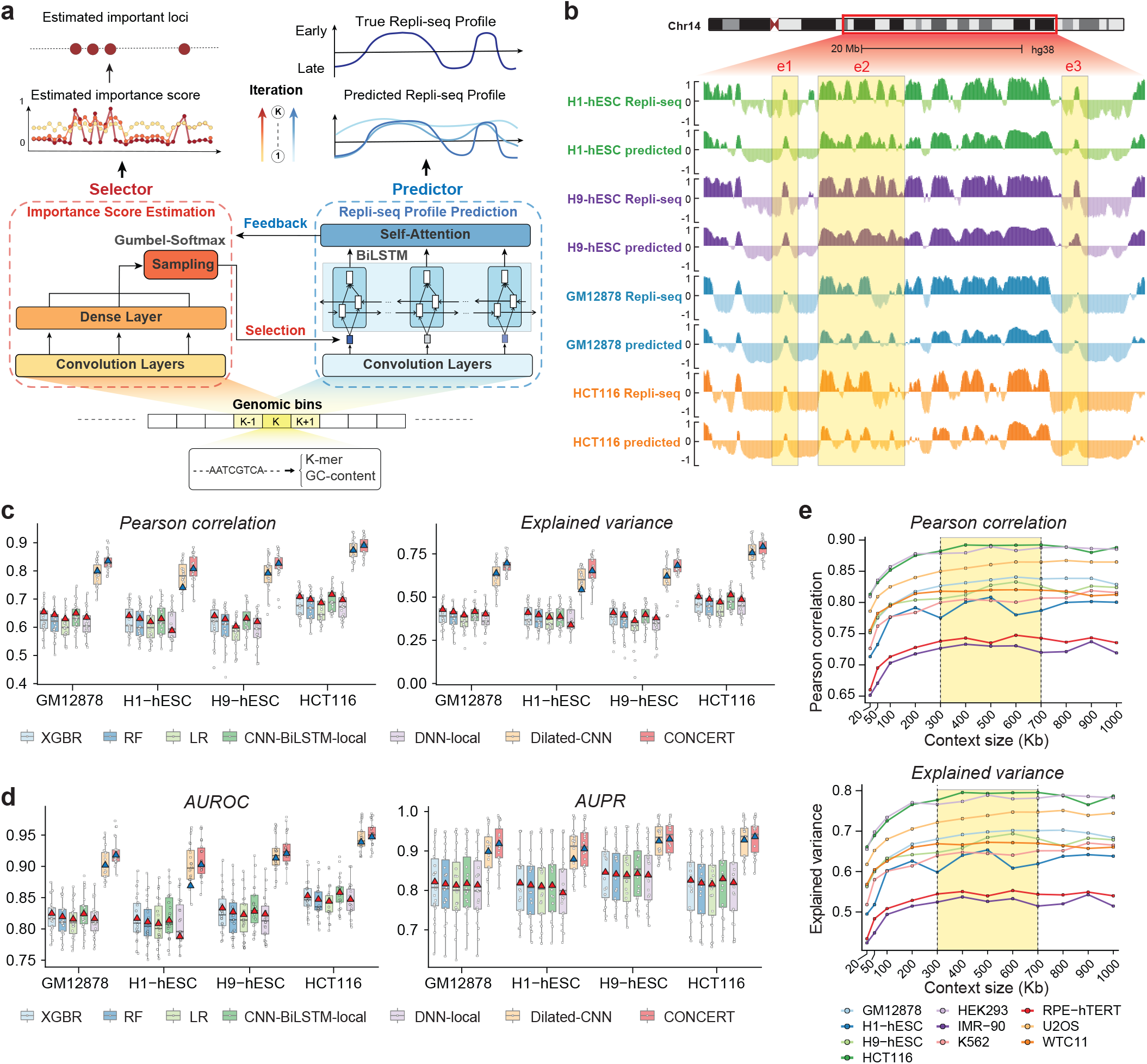
Overview of the Concert model and the performance evaluation in human cell types. **a**. Schematic of the Concert model framework. The detailed model architecture is shown in **Fig**. S1. **b**. An example of the predicted RT profiles in a genomic region on chromosome 14 in H1-hESC, H9-hESC, GM12878, and HCT116 in comparison with the real Repli-seq signals as shown in the UCSC Genome Browser. The regions e1, e2, and e3 are examples showing that the predictions of Concert capture both the similar RT patterns shared cross cell types and the cell type-specific patterns. **c**. Performance evaluation of RT predictions by different methods in H1-hESC, H9-hESC, GM12878, and HCT116. Each boxplot shows the performance variation across 22 autosomes for the corresponding method in the given cell line. The red/blue triangle represents the performance of genome-wide predictions based on merging the predictions across chromosomes. Detailed evaluation results in H1-hESC are shown in **Table** S3. **d**. RT classification performance across chromosomes in the four cell lines above. **e**. The change of RT prediction performance with respect to different context size of each genomic locus in 10 human cell lines. For **c** and **d**, performance evaluations in 10 human cell types are shown in **Fig**. S2 and **Fig**. S3.

### Concert achieves high RT prediction accuracy in multiple human cell types

To evaluate the performance of Concert, we carried out cross-chromosome RT prediction in 10 human cell types by using two subsets of chromosomes for model training (**Supplementary Methods** A.2.1; see **Fig**. 1b for examples), obtaining genome-wide RT predictions and importance score estimation. We included six other methods for performance comparisons: XGBoost Regression (XGBR) [23], Random Forest (RF), Linear Regression (LR), DNN-Local, CNN-BiLSTM-Local (adapted from DanQ [11]), and Dilated-CNN (adapted from the dilated convolution module of Basenji [18]) (**Supplementary Methods** A.2.3-A.2.4). We also adapted the full model of Basenji for RT prediction (**Supplementary Methods** A.2.5); however, we found that this adapted Basenji full model had lower performance than Dilated-CNN. We therefore only included Dilated-CNN in the performance benchmarking. We assessed the average and variation of prediction performances across different chromosomes in each cell type (**Fig**. 1c, **Fig**. S2, **Supplementary Methods** A.2.2). We also merged the predictions on all chromosomes to report genome-wide results (**Tables** S1 and S2). Overall, Concert consistently outperformed other methods in all cell types. For Concert, the median Pearson correlation coefficient (PCC) and median Spearman’s rank correlation coefficient (Spearman’s *ρ*) between the genome-wide predicted RT and the real RT signals across the 10 human cell lines are 0.81 and 0.80, respectively. In particular, Concert showed clear advantages over other methods that do not use context information. Notably, Concert outperformed Dilated-CNN in different cell types with respect to different evaluation metrics. This evaluation confirms the advantage of Concert that predicts RT profiles and identifies predictive RT-modulating genomic loci simultaneously.

The performance of Concert varies across cell types, with relatively high prediction accuracy in HCT116 and HEK293 (above 0.85 for both PCC and Spearman’s *ρ*) while lower accuracy in IMR-90 and RPE-hTERT, suggesting that there may exist cell type-specific dependencies between DNA sequence features and RT patterns. Concert has the capability of capturing not only the similar RT patterns shared by different cell types but also cell type-specific RT patterns (see examples in **Fig**. 1b). We further evaluated Concert using different context sizes (**Fig**. 1e). We found that the prediction accuracy increases as the context expands within a range in each cell type, indicating the informativeness of long-range context for RT prediction. The performance approaches saturation as the context size exceeds +/-150Kb. In addition, we evaluated our method on early or late RT classification (**Supplementary Methods** A.2.2). Concert reached 0.81-0.95 in AUROC (area under the receiver operating characteristic curve), 0.72-0.96 in AUPR (area under the precision-recall curve) in 10 human cell types, consistently outperforming the other methods (**Fig**. 1d, **Fig**. S3).

The numbers of predicted important genomic loci for RT in different human cell types that are also shared by the other cell types are listed in (**Table** S7). We also identified cell type-specific important loci (**Table** S8). The cross cell-type comparison suggests that there may exist sequence elements with conserved or cell type-specific functional importance in modulating RT across cell types. Specifically, we identified ESC-specific important genomic loci that are only predicted in H1-hESC or H9-hESC. Gene Ontology (GO) analysis of the identified important loci using GREAT [24] showed that they are significantly associated with fundamental developmental processes (**Fig**. S5). Also, based on the gene expressions in H1-hESC, GM12878, and K562, both cell type-specific expressed genes and genes with cell type-specific increased expressions are more enriched near the predicted important loci and show higher enrichment around the predicted cell type-specific important loci, both in early RT regions and genome-wide (**Fig**. S6, **Supplementary Methods** A.3.8).

Together, these results strongly demonstrate the effectiveness and advantages of Concert in learning functionally important genomic loci to predict RT profiles from DNA sequences.

### Concert predicted important loci in mESCs show consistency with ERCEs

Next, we applied Concert to predict RT in mESCs and evaluated the importance scores by comparing to the recently discovered ERCEs in mESCs [8]. In [8], multiple *cis*-regulatory elements with crucial roles in controlling early replication (i.e., ERCEs) were identified in mESCs and were validated by CRISPR-mediated genome engineering. In particular, three cooperative ERCEs were discovered at the *Dppa*2/4 domain on chromosome 16 in mESCs (denoted as ‘a’, ‘b’, and ‘c’ in **Fig**. S4) and the targeted deletion of these three elements caused a switch from early to late replication [8]. Additionally, 1,835 ERCEs were computationally inferred genome-wide based on specific epigenetic properties shared with the three experimentally identified ERCEs. Two ERCEs (‘d’ and ‘e’ in **Fig**. 2b) inferred at the *Zfp42/Rex1* locus on chromosome 8 were also further verified experimentally.

**Figure 2.**
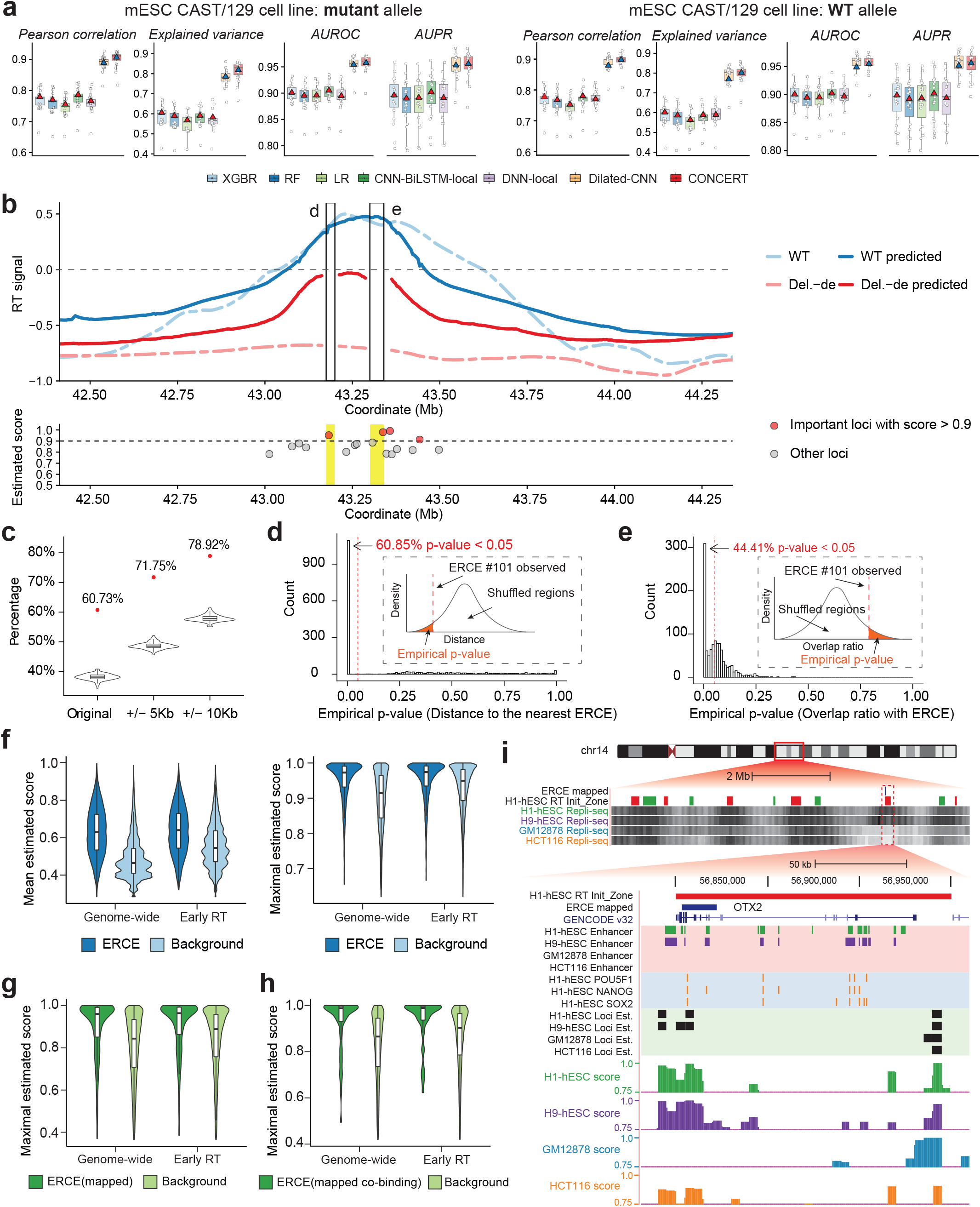
Evaluation of the predictions from Concert in mESCs and hESCs. **a**. RT prediction performance (left panel on each row) and early/late RT classification performance (right panel on each row) on the mutant and WT alleles of mESC CAST/129 in comparison with different methods (Repli- seq data from [8]). **b**. RT prediction with respect to the presence of ERCEs at the *Zfp42/Rex1* locus on chromosome 8 in mESCs and the identified important genomic loci with high Concert importance scores. Normalized estimated importance scores below 0.5 are not shown. **c**. Percentage of the predicted ERCEs (the red dots) that overlap with the Concert predicted important loci, with varying extension sizes of the original ERCEs. The boxplots show the corresponding background distributions of the percentage of the randomly sampled regions overlapping with the predicted important loci. **d**. Distribution of the empirical *P*-value of the distance of a predicted important locus to the nearest ERCE. The predicted loci are randomly shuffled on the corresponding chromosomes to estimate the empirical background distribution. The sub-plot included is an example showing the distance of a predicted important locus to the nearest ERCE vs. the background distribution of the distances for the randomly shuffled loci. **e**. Distribution of the empirical *P*-value of the overlap ratio of a predicted important locus with the matched ERCE. The predicted loci are randomly shuffled on the corresponding chromosomes to estimate the empirical background distribution. The sub-plot included is an example showing the overlap ratio of a predicted important locus with the matched ERCE vs. the background distribution of the overlap ratios for the randomly shuffled loci. **f**. Distribution of the estimated importance scores of ERCE regions vs. the background distribution in non-ERCE regions (left: region-wise mean scores; right: region-wise maximal scores). **g**. Distribution of the estimated importance scores of ERCE regions mapped to human vs. the background distribution in H1-hESC. **h**. Importance score distribution of ERCE regions mapped to human with the co-binding sites of POU5F1, SOX2, and NANOG in H1-hESC. **i**. Examples of the RT prediction and cell type-specific important genomic loci identified on chr14:54,295,000-58,750,000. Loci Est. represents the estimated RT-predictive important genomic loci. H1-hESC Init_Zone represents replication initiation zones in H1-hESC [27].

We sought to explore the connection between ERCEs and the important RT-modulating genomic loci identified by Concert using sequence features only. We used the same mESC Repli-seq data in CAST/129 hybrid cells from [8]. There are two alleles: mutant allele, with targeted ERCE deletions, and WT (wild type) allele, which share similar RT profiles except for the ERCE deleted regions. The similar model training procedures applied to RT prediction in the human cell types were applied to mESCs (**Supplementary Methods** A.2.1). Concert achieved high RT prediction accuracy in mESCs, with PCC 0.90, 0.91, explained variance 0.80, 0.79 on the mutant and WT alleles, respectively, outperforming other methods (**Fig**. 2a). We next identified RT-modulating genomic loci based on the estimated importance scores. Specifically, we normalized the scores to the scale [0,1] using quantile transformation followed by detecting local peaks of scores to select a subset of loci with distinguishable prominence (**Supplementary Methods** A.3.1). Concert identified seven important genomic loci at the *Dppa*2/4 locus on mouse chromosome 16, where three directly overlap with ERCEs ‘a’, ‘b’, ‘c’ and two overlap with ‘b’ and ‘c’ with +/- 1 bin (5Kb) extension (**Fig**. S4). Also, Concert identified three important genomic loci at the *Zfp42/Rex1* locus on chromosome 8 (**Fig**. 2b), where two loci directly overlap with ERCEs ‘d’ and ‘e’. Altogether, each of the five ERCEs that were experimentally validated by CRISPR-mediated deletions in [8] was matched by RT-modulating genomic loci predicted from Concert. Concert also predicted the local switch from early to later RT after the *in silico* deletion of each of the two sets of ERCEs (‘a’, ‘b’, ‘c’ and ‘d’, ‘e’), consistent with the experimental results in [8] (**Fig**. 2b and **Fig**. S4).

We next compared the identified RT-modulating genomic loci genome-wide and the predicted ERCEs from [8] (1,798 predicted ERCEs on autosomes with 20Kb∼200Kb in length) We found that 1,419 of the 1,798 predicted ERCEs with +/-2 bin extension (78.92%, empirical *P<*1e-02, **Fig**. 2c) overlap with one or more predicted RT-modulating loci identified by Concert. The Concert predicted impor- tant loci also have shorter distance to the nearest ERCEs and higher overlap with the matched ERCEs as compared to the background where the elements are randomly shuffled (**Fig**. 2d-e, *P<*2.2e-16 with Kolmogorov-Smirnov test, or K-S test [25]). We further evaluated the Concert estimated importance scores for ERCEs compared with the randomly sampled non-ERCE sequences (background). ERCEs on average have significantly higher region-wise maximal and mean scores from Concert than the back- ground, both in early RT domains and genome-wide (**Fig**. 2f, *P<*2.2e-16 with K-S test, **Supplementary Methods** A.3.2).

Together, these results further confirm the effectiveness of Concert in predicting genome-wide important RT-modulating sequence elements supported by ERCEs in mESCs.

### Concert predicted important loci in human ESCs match orthologous ERCEs

As the ERCEs were only inferred in mESCs [8], we aimed to assess how conserved these important *cis*-elements for RT are between human and mouse in terms of their connection to the RT-modulating elements predicted by Concert. By mapping the mouse ERCEs to the human genome with reciprocal liftOver [26] (**Supplementary Methods** A.3.2), we obtained 446 mapped ERCEs and evaluated the Concert importance scores of the mapped regions in H1-hESC. We compared the distribution of region-wise maximal (or mean) estimated importance scores of the mapped ERCEs with the background distribution based on randomly sampled regions in H1-hESC (**Fig**. 2g, **Supplementary Methods** A.3.2). The mapped ERCEs in human exhibit higher Concert importance scores on average compared to the background in both early RT domains and genome-wide (*P<*2.2e-16). The mapped ERCEs in human with co-binding of two or more pluripotency TFs also show higher Concert importance scores than the background (*P<*1e-7, **Fig**. 2h). Furthermore, 71.08% of the mapped ERCEs in human with +/-2 bin extension can be matched by Concert identified RT-modulating loci (empirical *P<*1e-02).

An example of the mouse ERCE mapped to the human genome that can be identified by Concert is shown in **Fig**. 2i, This is a genomic region with hESC-specific early RT and contains a co-binding of the pluripotency TFs POU5F1, SOX2, and NANOG in H1-hESC. The mapped ERCE also resides in a replication initiation zone in H1-hESC identified from [27] (**Fig**. 2i). Concert identified cell type-specific RT-modulating loci that overlap with the mapped ERCE in hESC (supported by data in both H1-hESC and H9-hESC). The non-ES cell types such as GM12878 and HCT116 exhibit late RT locally, without important genomic loci identified by Concert at the same location. Also, the 5’ side of the TF-encoding gene OTX2 is co-localized with the identified important loci in H1-hESC and H9-hESC, with hESC-specific expression (**Supplementary Methods** A.3.8). OTX2 has critical roles in the early stages of embryonic development and ESC differentiation [28, 29]. There are cell type-specific enhancers (annotations from [30]) near the promoter of OTX2 in H1-hESC and H9-hESC, which overlap with the pluripotency TF co-binding site and the Concert identified important loci. This points to the potential connections underlying the OTX2 regulation, the co-binding of pluripotency TFs, and the ESC-specific early RT modulation at this locus. This mapped ERCE in human is highly likely to be a human ES cell-specific ERCE and there is potential functional conservation between human and mouse (**Fig**. 2i).

These analyses provide additional evidence regarding the potential of Concert in identifying RT- modulating sequence elements in human cell types.

### Concert predicted RT-modulating loci are enriched with specific regulatory elements

To explore the connection between transcriptional regulatory elements and the Concert predicted RT- modulating sequences, we compared the estimated importance scores and the annotated *cis*-regulatory elements (CREs) in different human cell types. The CRE annotations were retrieved from the SCREEN database [31], comprising mainly five types of CREs: dELS (distal enhancer-like signatures), pELS (proximal enhancer-like signatures), PLS (promoter-like signatures), DNase-H3K4me3 (likely poised elements with DNase and H3K4me3 marks), and CTCF-only. We found that the loci with pELS, PLS, or DNase-H3K4me3 signatures received significantly higher estimated importance scores from Concert than the non-CRE loci, which was consistently observed in different cell types (**Fig**. 3a, *P<*2.2e-16, **Supplementary Methods** A.3.3).

We further analyzed the TF binding enrichment in the Concert predicted important loci in H1- hESC using the ChIP-seq data from ENCODE [32] (**Supplementary Methods** A.3.3). We observed enriched presence of specific TF binding sites in the Concert predicted important loci as compared to the background loci with matched RT signal levels (**Fig**. 3c,**Supplementary Methods** A.3.4), including the TFs known to be critical for transcription or the pluripotency of ESCs. Specifically, the binding sites of the pluripotency TFs, POU5F1, SOX2, and NANOG, are significantly enriched in the Concert predicted important loci (Fisher’s exact test, *P<*0.01), of which POU5F1 in particular shows high enrichment (*P<*2.2e-16). These three TFs were also identified as key signatures of mESC ERCEs in [8].

We further compared the chromatin states annotated by ChromHMM [33] in the Concert predicted RT-modulating loci in different human cell types (**Fig**. 3b). We merged the 15 chromatin states into 8 groups using the definition in [34] (**Supplementary Methods** A.3.5). We observed distinct patterns of the different chromatin state groups in the predicted important loci.

**Figure 3:**
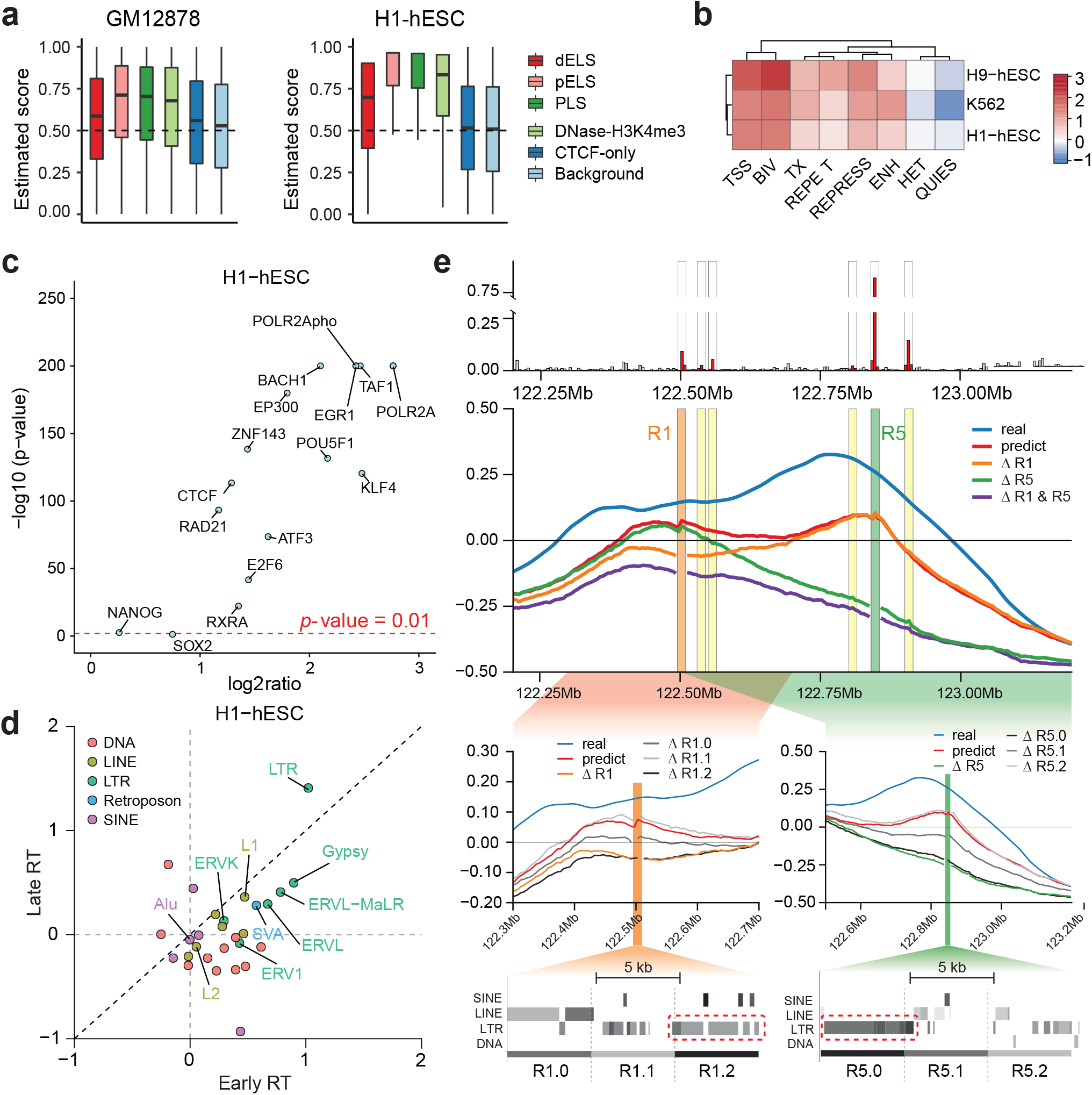
Comparisons between the Concert predicted important loci and different genomic and epigenomic features. **a**. Distribution of importance scores in genomic loci with CREs in open chromatin regions in GM12878 and H1-hESC. **b**. Enrichment of chromatin states annotated by ChromHMM [33] in the predicted important loci. **c**. Enrichment of specific TF or non-TF binding sites in the predicted important loci in open chromatin regions in H1-hESC. The enrichment is measured by log2 Fisher’s exact test statistics (x-axis) and the *P*-value (y-axis). **d**. Log2 fold change of the percentage of predicted important loci depleted of known regulatory elements and overlapping with a specific TE family vs. the percentage of predicted loci overlapping with both regulatory elements and the specific TE family in early and late RT regions in H1-hESC. **e**. Examples of predicted important loci (depleted of regulatory elements) with specific TE families on chr6:122,250,000-123,000,000 in H1-hESC. 1st panel: the estimated importance scores; 2nd panel: real RT and predicted RT without or with deletions; 3rd panel: predicted RT with deletion per bin for R1 or R5; 4th panel: TE families in each bin of R1 or R5. The important bins and associated TEs are marked with dashed lines.

Together, these results suggest that the predicted RT-modulating sequence elements from Concert co-localize with specific types of transcriptional regulatory elements, indicating the intertwined nature of transcription and DNA replication.

### Specific transposable elements are over-represented in the Concert predicted impor- tant loci depleted of regulatory elements

Our analyses revealed that the predicted important loci for RT show signatures of specific transcription-related regulatory elements. However, we have limited knowledge of what sequence features may exist in the DNA elements with RT-modulating functions beyond the known regulatory elements. We next specifically explored the connections between the transposable elements (TEs)/repetitive elements (REs) and the estimated important loci globally, using the annotations in the human genome from the UCSC Genome Browser [35] (**Supplementary Methods** A.3.6). We observed specific types of repetitive sequences are enriched in genomic loci with high importance scores from Concert (**Figs**. S7-S8). We found that Alu elements show strong preference in loci with high estimated importance in different cell types, consistent with earlier studies [1]. Additionally, scRNA, snRNA, srpRNA, and SVA are also highly represented in loci with high Concert estimated scores, in accordance with findings from previous studies on RT comparison across multiple primate species [5].

Next, we analyzed the features of the predicted important loci that are depleted of known regulatory elements. We used multiple annotations to identify the genomic loci without regulatory elements in human cell types, including chromatin states, CREs, and enhancers (**Supplementary Methods** A.3.7). Notably, the Concert predicted important loci depleted of regulatory elements in different human cell types show enrichment of specific transposable element (TE) families or REs as compared to the predicted important loci overlapping with regulatory elements (**Fig**. 3d). In H1-hESC, the corresponding enriched repetitive sequences include multiple TE families of LTR (including LTR, ERVL, ERVL-MaLR, ERVK, ERV1, Gypsy), L1 elements, and SVA. The enrichment patterns were observed in both early and late RT regions, and consistent across cell types (**Fig**. S9).

As an example, there are 6 predicted important loci depleted of regulatory elements in a genomic region displaying local early RT on chromosome 6 in H1-hESC (**Fig**. 3e). Concert predicted the emergence of the local early RT. We performed *in silico* deletions of these 6 loci (denoted as R1-R6, with +/-1 bin extension each) both individually and in combinations to evaluate the effect of each locus or the combinations. By deleting both R1 and R5, the local predicted RT switched from early to late, exhibiting the most significant change among all the deletions. Deleting R1 or R5 alone changed the corresponding side of the RT profile from early to late, while the other side of the RT profile remained early RT. Deleting R5 alone induced the strongest prediction change among the individual deletions. Note that Concert predicted the highest importance score at R5 among these 6 loci. Deleting both R1 and R5 resulted in larger change of predicted RT than the accumulation of predicted RT decrease by individual deletions, suggesting the potential synergistic cooperative functions of R1 and R5. We then searched for the distinctive sequence features underlying the two loci. As R1 and R5 each contain +/-1 bin extension (3 bins in size, denoted as R1.0-R1.2 and R5.0-R5.2, respectively), we deleted each bin separately *in silico* and compared the change of predicted RT (**Fig**. 3e). The loci with the strongest effects (R1.2, R5.0, the deletion of which is similar to deleting the whole R1 or R5) are each primarily composed of LTR elements; specifically, ERVK (R1.2) and ERV1 (R5.0), consistent with the observations in **Fig**. 3d. Moreover, based on the multi-species sequence alignment, the loci with strong effects correspond to sequences only present in human and the closely-related primates (**Fig**. S10). This strongly suggests that the TEs in the two loci were likely inserted into the genome during recent primate evolution, which may be subsequently exapted to perform lineage-specific function of RT modulation.

This analysis suggests the existence of a potential new class of RT-modulating elements that are depleted of known transcriptional regulatory elements but are strongly represented by specific TEs/REs.

### The interplay between the Concert predicted important loci and replication origins as well as 3D chromatin interactions

We next asked if our Concert predicted important loci co-localize with the replication origins identified from the SNS-seq data [36]. We performed peak calling to locate RT origins using the SNS-seq data in H1-hESC and H9-hESC cell lines from [27]. The genomic loci overlapping with RT origins have on average significantly higher estimated importance scores than the loci without identified RT origins (**Fig**. 4a, *P<*2.2e-16, K-S test, **Fig**. S11). In particular, 13,230 of the predicted important loci (33.54%) overlap with a SNS-seq derived RT origin in H1-hESC (**Fig**. 4b). This suggests that Concert identifies a subset of the RT origins. The identified non-RT origin but important genomic loci from Concert may harbor sequence elements that participate in RT regulation but not RT origins. We also found that the RT origins exhibit discrepancy of estimated important scores. The importance scores estimated by our method may reflect the heterogeneity of the RT origins (see examples in **Fig**. 4f-g).

**Figure 4:**
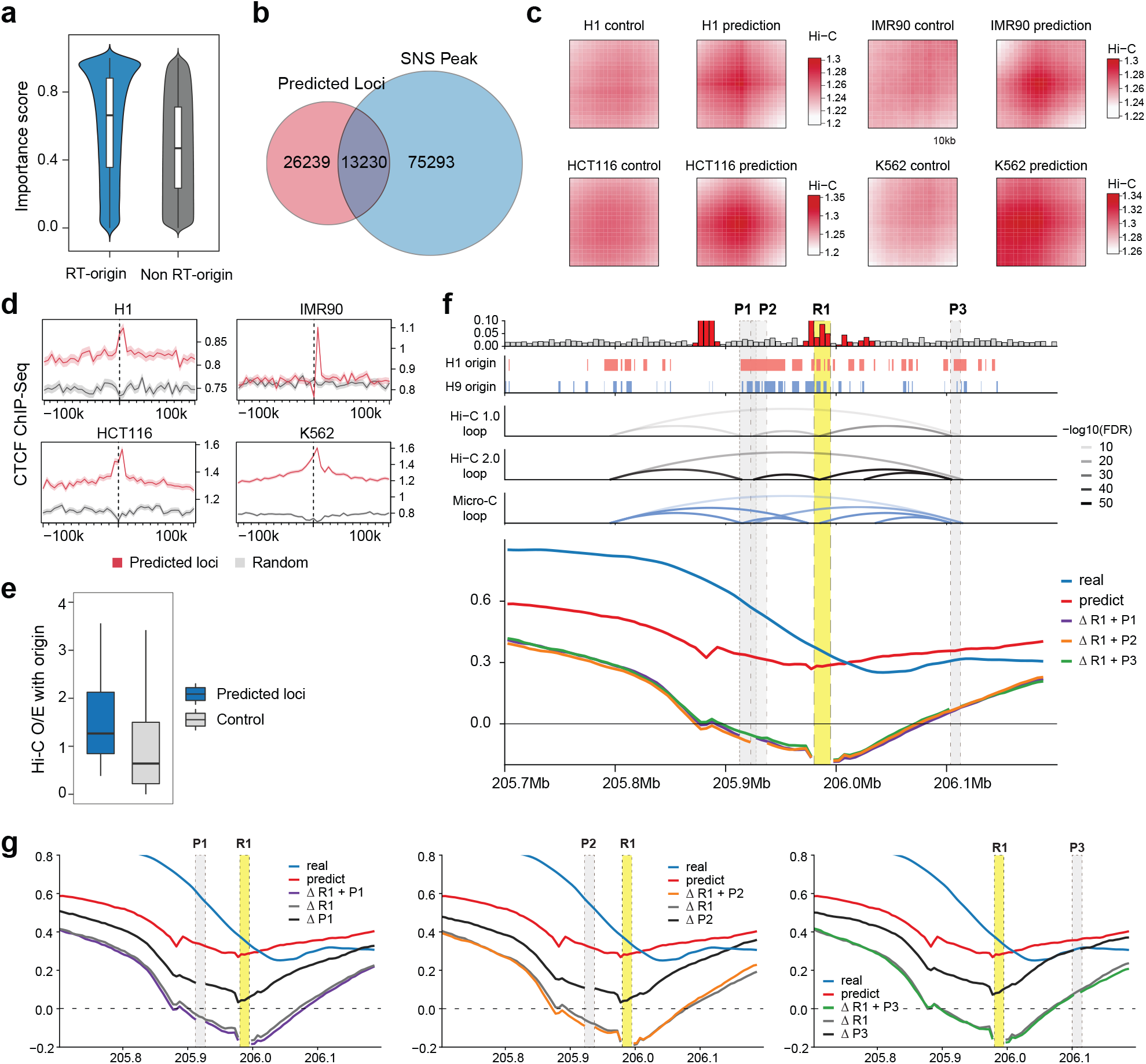
Interplay between the Concert predicted RT-modulating important loci and RT origins as well as 3D chromatin interactions. **a**. Distribution of the estimated importance scores on RT origins and non-origins. RT origins were defined by SNS-seq peaks in H9-hESC [27]. **b**. The overlap between the predicted important loci and SNS-seq peaks in H9-hESC. **c**. Hi-C aggregate analysis of the predicted important genomic loci. **d**. Enrichment of CTCF ChIP-Seq scores in genomic regions surrounding the predicted important loci. **e**. Distribution of Hi-C O/E scores of interactions between the predicted important loci and RT origins. **f**. Genome Browser example of the predicted important loci in the context of 3D chromatin interactions in H1- hESC. The tracks shown are (from top to bottom): estimated importance scores, RT origins identified from SNS-seq data (H1-hESC and H9-hESC) [27], Hi-C loops, Micro-C loops [59], real RT and predicted RT with or without deletions of specific pairs of genomic loci. **g**. Predicted RT with individual *in silico* deletions of the genomic locus in each specific pair of loci. Predicted RT with paired deletions is also shown.

RT is highly interconnected with 3D genome organization. To further reveal the potential dependencies between the estimated RT-modulating loci and the 3D genome features, we performed Hi-C aggregate analysis of the predicted important loci in different human cell types (**Supplementary Methods** A.3.9). The identified important genomic loci showed higher proportion of Hi-C contacts compared to the background in different cell types (**Fig**. 4c). As CTCF is known to be crucial for mediating chromatin loops, we analyzed the binding distribution of CTCF in genomic regions harboring the Concert predicted important loci. We found that the Concert predicted important loci have higher CTCF binding scores as compared to the surrounding regions, while the enrichment pattern was not found in the control group (**Fig**. 4d, **Supplementary Methods** A.3.9).

Next, we assessed the potential chromatin interactions between the Concert predicted important loci and the RT origins identified from the SNS-seq data [27], by which the Concert predicted loci interact with RT origins to cooperatively modulate RT. We calculated the Hi-C O/E (observed/expected) interaction score for each pair of predicted important genomic locus and an RT origin within a distance range. The predicted important loci show significantly higher average Hi-C O/E scores than the control (**Fig**. 4e, *P<*2.2e-16, K-S test, **Supplementary Methods** A.3.9), indicating that the Concert predicted RT-modulating loci show spatial interactions with the RT origins.

As an example, we show the connections between the predicted important loci and the RT origins in the context of 3D genome organization (**Fig**. 4f-g). Here, Concert predicted an important genomic locus for RT (denoted as R1) in an early RT region on chromosome 1 in H1-hESC, which overlaps with the SNS-seq derived RT origins and is connected to three RT origins (P1-P3) by two chromatin loops (**Fig**. 4f, **Supplementary Methods** A.3.9). R1 is among the loci with the highest estimated importance scores in the region, while P1-P3 receive lower scores. To evaluate the importance of R1 and P1-P3, we performed individual and paired *in silico* deletions of the loci (**Fig**. 4f-g). Each pair involves R1 and one of P1-P3 connected by a Hi-C loop. We found that each of the paired deletions caused the local predicted RT to switch from early to late (**Fig**. 4g). For individual deletions, deleting R1 caused the most significant effect, showing the change of early to late RT. Deleting each of P1-P3 alone caused moderate decrease of the predicted RT signals. These observations suggest that R1 is a strong candidate locus for RT modulation. Importantly, the Hi-C loops connecting R1 to the other RT origins may assist R1 to synergistically regulate RT.

In summary, these results demonstrate that Concert helps unveil the interplay among the RT- modulating sequence elements, the replication origins, and 3D chromatin interactions.

## Discussion

In this work, we developed Concert to predict DNA replication timing profiles using DNA sequence features only, with the model training procedure using the Repli-seq data as input. Application to 10 human cell lines demonstrated that Concert reached high prediction performance in a wide range of cell types and outperformed other methods. The results additionally showed conservation and variation in the potential dependencies between DNA sequences and RT signals in different cell types. Our method also reached high RT prediction accuracy in mESCs. For predictive sequence element identification in mESCs, each of the five ERCEs in mESCs that were experimentally validated through CRISPR-mediated deletions in [8] can be identified by our method. We also observed connections between the Concert predicted RT-modulating elements and different types of genomic, epigenomic, and 3D genome features, unveiling sequence elements that may have underappreciated roles for RT modulation. Our method demonstrates the potential for prioritizing experimental characterizations of possible sequence determinants of RT. Overall, Concert provides a generic framework to predict large-scale genomic features and discover the underlying predictive sequence elements that may modulate RT program using DNA sequence features only.

There are a number of possible improvements and future directions for Concert. First, the model can incorporate specific cooperative functions of sequence elements, as revealed in [8], where several ERCEs were found to cooperate as a group to regulate early RT in the local domain. Second, we can include other types of epigenetic features into the model to reflect cell type-specific characteristics of RT regulation, and improve the model performance in RT prediction and recognition of possible RT regulators. We can also utilize the integration of epigenetic features to perform cross cell type RT predictions. Finally, to further disentangle sequence determinants for RT, transcription, and 3D chromatin structures that are intrinsically interconnected, an integration of Concert and other related sequence-based models [19, 22] may offer a systematic solution to connect genomic sequence, genome structure, and genome functions. For example, we found the Concert predicted important loci have stronger CTCF binding than their flanking regions. However, earlier work showed that CTCF depletion does not lead to changes in genome-wide RT program in mESC [8]. Future work is needed to clearly demonstrate the mechanistic connections between Concert predicted RT-modulating loci and chromatin folding. Nevertheless, we have demonstrated that Concert has shown strong promise to help uncover important sequence level properties that potentially modulate the genome-wide RT program in a diverse set of cell types. The Concert predictive model also has the potential to help expand the catalog of functionally-relevant genetic variants in the human genome.

## Methods

### Concert – Interpretable RT prediction using sequence features with context

In Concert, we aim to maximize the mutual information between the target genomic signals (RT in this work) to be predicted and the underlying DNA sequences, with a set of predictive sequences selected. We divide the genome into non-overlapping consecutive bins of equal sizes. Each genomic bin is a locus. Suppose we use +/- *l* bins around each locus to form a context. The context surrounding each locus forms a sliding window of size *L* = 2*l* +1 along the genome. Features of loci in each context window are fed to the model to predict locus-wise signals. Suppose *f*_*θ*_(*x*) is the function of the predictor, which maps the sequence features *x* ∈ ℝ^*L*×*d*^ to the signals *y* ∈ ℝ^*L*×*c*^, with *θ* as the parameters, and *d, c* representing the sequence feature dimensionality and the number of genomic signal types, respectively. We use *c* = 1. Our model is extendable to predict multiple types of genomic signals with *c* > 1.

Let *X* = (*X*_1_, …, *X*_*L*_), *Y* = (*Y*_1_, …, *Y*_*L*_), where *X*_*k*_ ∈ ℝ^*d*^, *Y*_*k*_ ∈ ℝ denote the feature vector and the RT signal of the *k*-th genomic locus, respectively, *k* = 1, …, *L*. Let ***𝒱***^(*L*)^ = {*ν* ∈ ℝ^*L*^|*ν*_*k*_ ∈ {0, 1}, *k* = 1, …, *L*}. Let *S* = (*S*_1_, …, *S*_*L*_) ∈ ***𝒱***^(*L*)^ denote a specific combinatorial selection of predictive loci, with *S*_*k*_ = 1 indicating the *k*-th locus is selected and *S*_*k*_ = 0 otherwise. Suppose *ϕ*_*β*_(·) is the function of the selector, mapping *X* to *S*, with *β* as the parameters. Let *X*_*S*_ be the representation of *X* with only the features of selected genomic loci retained. The original objective function is (more details are in **Supplementary Methods** A.1.1):

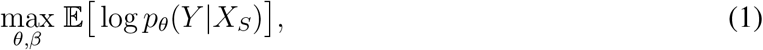

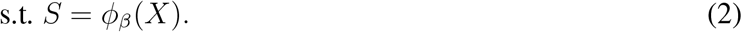

#### The selector module in Concert

With the selector module we aim to select a subset of genomic loci that are predictive of the RT signals in each context window. Let 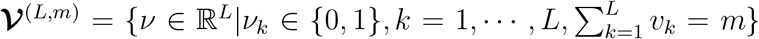 denote all the possible selections of *m* loci from *L* loci, 1 ≤ *m* ≤ *L*. Suppose we first only select one predictive locus from the window. Suppose the selection for a given context window is denoted by 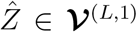.. Let *z* = (*z*_1_, *z*_2_, …, *z*_*L*_) ∈ ℝ^*L*^ be the category probabilities of locus selection, where each category corresponds to a locus. We use the Gumbel-Softmax reparameterization approach [37] to approximate the distribution of 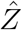 with a Concrete distribution which is differentiable.

First, we add randomness to the probability of selecting each locus using random variables 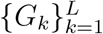 that are independently sampled from a Gumbel distribution [38]. Let *G*_*k*_ = − log(− log *U*_*k*_), *U*_*k*_ ∼ Uniform(0, 1), where Uniform(0, 1) represents uniform distribution on [0, 1]. We have *G*_*k*_ ∼ Gumbel(0, 1), *k* = 1, …, *L*. The random perturbation facilitates learning robust selection that is less sensitive to noise.

Suppose *α*_1_, …, *α*_*L*_ are unnormalized category probabilities, *α*_*k*_ > 0. If we use the Gumbel-Max approach [37], we select the 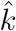-th locus as the predictive locus with the objective function:

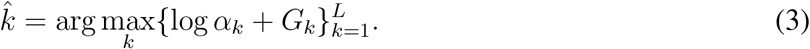

The objective function is not differentiable everywhere in its domain. We employ the Gumbel-Softmax reparameterization trick that was developed for the continuous relaxation of the arg max function [37]. Using the Gumbel-Softmax trick, the probability of selecting locus *k* as the predictive locus is:

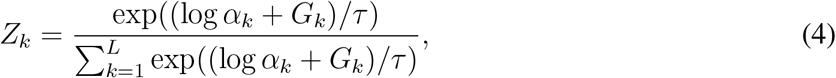

where *τ* is a temperature parameter. The softmax function approaches the arg max function as *τ* approaches zero, with the orders of {log *α*_*k*_ + *G*_*k*_} preserved [37]. We denote the distribution of *Z* = (*Z*_1_, …, *Z*_*L*_) ∈ ℝ^*L*^ as a Concrete distribution parameterized by (*α, τ*), where *α* = (*α*_1_, …, *α*_*L*_). We have *Z* ∼ Concrete(*α, τ*).

To sample a subset of *m* predictive genomic loci from the *L* loci, we sample a vector *Ŝ* from **𝒱** ^(*L,m*)^. We adopt the strategy proposed in [39] for the approximation of sampling *Ŝ*. First, we sample *m* Concrete random vectors *z*^(1)^, …, *z*^(*m*)^ independently from the distribution Concrete(*α, τ*). Second, for each genomic locus *k*, we take the maximal value at the *k*-th dimension of the sampled random vectors as the estimated probability of selecting locus *k*. Let 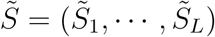, where 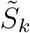 represents the probability of selecting locus *k*. 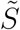 is a continuous approximation of the discrete vector *Ŝ*. We have:

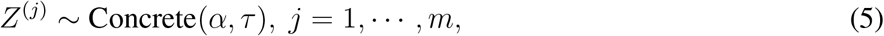

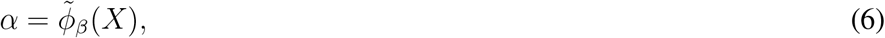

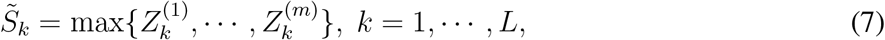

where 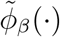is the function mapping the sequence features to *α* = (*α*_1_, …, *α*_*L*_) in the selector module, with parameters *β*. We interpret *α*_*k*_ as the estimated importance score of the *k*-th locus.

Let *G* denote the set of random variables 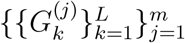 which are sampled from the Gumbel distribution. We havep 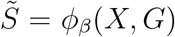. Suppose the number of genomic loci is *N*. As each locus is the center of a context window, the number of context windows is *N*. With regularization on the model complexity to reduce model over-fitting, the objective function is:

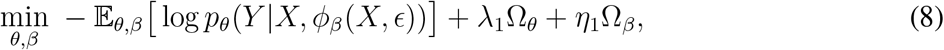

Where 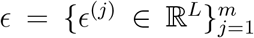 consists of random variables independently sampled for *x* ∈ ℝ^*L*×*d*^. We have 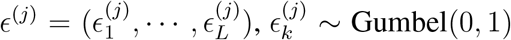, *k* = 1, …, *L, j* = 1, …, *m*. Ω_*θ*_, Ω_*β*_ represent model complexities of the predictor and the selector, respectively, with *λ*_1_, *η*_1_ as the corresponding regular- ization coefficients. We use *l*_2_-norm of parameters for measuring Ω_*θ*_ and Ω_*β*_ in practice. In the implementation, the selector comprises one dense layer for preliminary feature transformation, a convolution layer to utilize limited local context information for feature representation, followed by dense layers to generate estimates of *α*, and a Gumbel distribution based random variable sampler (**Supplementary Methods** A.1.3).

#### The predictor module in Concert

The predictor module consists of four sequentially connected parts (**Supplementary Methods** A.1.2). The first part is a preliminary feature transformation sub-module which generates an intermediate feature representation for each locus, with an interface from the selector (**Fig**. S1). The output of the sub-module is weighted by the importance scores *s*_*i*_ = *ϕ*_*β*_(*x*_*i*_, *ϵ*_*i*_) = (*s*_*i*,1_, …, *s*_*i,L*_) ∈ ℝ^*L*^ estimated by the selector, where *i* represents the index of the context window. The weighted feature is 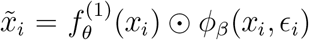, where 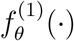 is the mapping function of the sub-module and ⊙ represents element-wise multiplication.

The second part is a BiLSTM layer [40], which enables bidirectional context information sharing across genomic loci (**Supplementary Methods** A.1.2). RNNs have been widely used for modeling sequential data [41–43]. LSTM and GRU [44, 45] are two representative variants of RNNs, using cell memory and gate mechanisms to learn long-range dependencies, with the information flow restricted to one direction. BiLSTM overcomes this limitation by splitting the hidden states of LSTM into forward and backward states, forming two parallel LSTM layers in different directions. The output of BiLSTM is the concatenation of the forward and backward states, producing context-aware feature representations. The third part is the self-attention layer, which is used to capture dependencies between two arbitrary loci in a context window [46]. With BiLSTM, information sharing is constrained by the spatial order of the loci. The shared information vanishes as the distance between loci increases, making it still challenging to model dependencies between distant loci. For each target locus *l* (represented by *x*_*l*_), the self-attention layer estimates an attention value of each locus *l*^′^ (including the target locus) in the context window, measuring the importance of *x*_*l*_*′* for predicting the signal of *x*_*l*_*′*. The features of the loci are weighted by their target-specific attention values and averaged to generate an updated context-aware feature vector of the target locus. The self-attention mechanism used is:

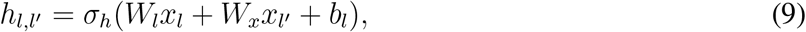

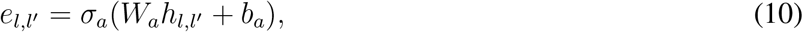

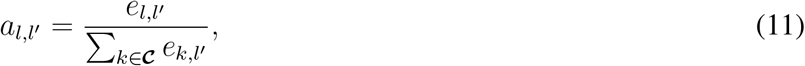

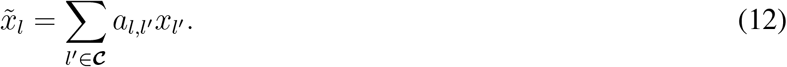

where ***𝒞*** represents the context window. *σ*_*h*_, *σ*_*a*_ denote nonlinear activation functions. {*W*_*l*_, *W*_*x*_, *b*_*l*_, *W*_*a*_, *b*_*a*_} are model parameters. The self-attention layer selectively aggregates the outputs from the BiLSTM layer to model distance-insensitive spatial dependencies across genomic loci. The fourth part is a fully-connected layer, mapping the features of each locus to a continuous value as the predicted RT signal. We use Keras/Tensorflow [47, 48] for implementation. The architecture details and hyperparameters of the model used in the analysis are described in **Supplementary Methods** A.1.3.

### Feature representation of genomic loci

#### Feature engineering with specific sequence patterns

We combine different types of features from each genomic locus as the input to the model, including the transformed *K*-mer frequency features, the GC profile based features, the optional transcription factor (TF) binding motif features, and the optional conservation score features (**Supplementary Methods** A.1.4). Let 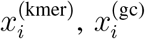 be the first two types of features of the *i*-th genomic locus, respectively. *K*-mer is a DNA subsequence of length *K*. The *K*-mer frequency feature is based on the occurrence frequency of each *K*-mer in the sequence of each locus, normalized by the sequence length, generating a 4^*K*^-dimensional feature vector. Let 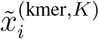 denote the feature for a specific *K*. For the balance between prediction performance and computation cost, we choose *K* = 5 and *K* = 6 to form concatenated fea- ture vector 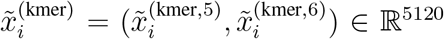. We choose PCA (Principal Component Analysis) for feature dimension reduction, with the reduced dimension *d*_*k*_ = 50. As GC content is known to correlate with RT [49], we extract two types of GC-based features, GC content (*f* ^(gc)^) and GC skew (*f* ^(skew)^). We have 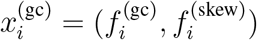.

#### Feature representation learning from local geomic sequences

We also propose incorporating a local-level sub-model which combines convolution neural networks (CNN) with BiLSTM to learn features from sequences (**Supplementary Methods** A.1.5), as an alternative to the pre-engineered feature representations. Integrating the CNN-BiLSTM sub-model, the model is re-formulated into a framework with a two-level hierarchical structure (denoted as Concert - hierarchical), providing a local-global multi-resolution scheme. At the first level, the sub-model learns to capture spatial dependencies within the sequence of each locus, functioning as a local feature extraction component. At the second level, the model learns dependencies across different genomic loci over larger-scale domains. The hierarchical structure shows similar RT prediction accuracy to the basic Concert model (**Table** S6). We therefore focus on the basic Concert model in the analysis in this work. Still, Concert -hierarchical provides an option to enable the flexibility of learning long-range spatial dependencies while remaining attentive to local sequence features.

### Data collection and processing

We used genome-wide RT maps based on Repli-seq data [50] in 10 human cell lines H1-hESC, H9-hESC, GM12878, HCT116, HEK293, K562, IMR-90, RPE-hTERT, U2OS, and WTC11. The two-fraction E/L Repli-seq data were downloaded from the 4DN Data Portal [51] and the ENCODE project [32] (accession numbers in **Tables** S4). We divided the genomes into consecutive non-overlapping 5Kb bins (the genomic loci). We followed the scripts on https://github.com/4dn-dcic/repli-seq-pipeline for Repli-seq analysis. We performed quality control of the Repli-seq reads using FastQC [52] and removed adapter sequences using Cutadapt [53]. We calculated the RT signal values for each non-overlapping 5Kb bin of the human genome. Specifically, we first mapped the processed sequencing reads to the human genome assemblies of hg38, using BWA [54]. The genome assemblies were downloaded from the UCSC Genome Browser [55]. We then calculated Repli-seq read count within a given genomic window (a genomic bin) in early and late phases of RT, respectively, normalized by the total mapped read count in early or late RT phase on the whole genome accordingly. The RT signal in each bin is defined as the base 2 logarithm ratio of read count per million reads between the early and late phases of RT within this region. Lastly, the log 2 ratio signal was quantile normalized and smoothed using the loess-smooth method [56].

The Repli-seq data in mouse were from the CAS/129 hybrid mESCs [8] (downloaded from GEO: GSE114139), which were derived from the cross between *M. casteneus* (CAST/Ei) and *M. musculus* (129/sv). SNP calls for CAST/Ei and 129/sv genomes were downloaded from: ftp://ftp-mouse.sanger.ac.uk/REL-1505-SNPs_Indels/. We first used Bowtie 2 [57] to map reads to a mouse genome (mm10) where all SNP positions are masked by the ambiguity base ‘N’. Then we used SNPsplit [58] to parse the mapped reads to the corresponding allele. Reads that do not overlap with SNPs were discarded. Reads overlapping with SNPs in the two genomes were sorted and processed in each allele individually. We followed the same scripts (https://github.com/4dn-dcic/repli-seq-pipeline) used in human RT analysis. The mouse RT signals in each allele were calculated at 5Kb resolution.

## Code Availability

The source code of Concert can be accessed at: https://github.com/ma-compbio/CONCERT.

## Acknowledgements

The authors would like to thank David Gilbert for sharing the Repli-seq data before publication and for feedback that improved the work. The authors are also grateful to Takayo Sasaki and Peiyao Zhao for their help with SNS-seq and Repli-seq data preparation. This work was supported by the National Institutes of Health Common Fund 4D Nucleome Program grant UM1HG011593 (J.M.) and the National Institutes of Health grants R01HG007352 (J.M.) and R01HG012303 (J.M.). J.M. is additionally supported by a Guggenheim Fellowship from the John Simon Guggenheim Memorial Foundation.

## Author Contributions

Conceptualization, Y.Y. and J.M.; Methodology, Y.Y. and J.M.; Software, Y.Y.; Investigation, Y.Y., Y.Z., Y.W., and J.M.; Writing – Original Draft, Y.Y. and J.M.; Writing – Review & Editing, Y.Y. and J.M.; Funding Acquisition, J.M.

