## Supplemental Information for "Concert: Genome-wide prediction of sequence elements that modulate DNA replication timing"

#### A Supplementary Methods

##### A.1 Details of the CONCERT model

###### *A.1.1 Overall framework of* CONCERT

We aim to maximize the mutual information between the selected input sequences and the corresponding RT signals, adopting the model interpretation mechanism introduced in [1]. In [1], instance-wise model explanation is performed as a separate step given a trained model. Specifically, two rounds of model training are performed, as firstly a task-specific predictive model is trained and secondly an explainer model is trained to interpret the predictive model. The training of the predictive model in the first step does not involve the explainer.

In CONCERT, we have made an important step to integrate a selector module with a predictor module to simultaneously predict the genomic signals and estimate the context-aware importance of each genomic locus for the prediction. The two modules share information between each other. The selector aims to select the predictive genomic loci in a given region and also assists the predictor in improving the prediction accuracy. Training is performed in one round, with the parameters of the selector and the predictor learned jointly.

Given the context window of size $L$ centered at a genomic locus, let $X = (X_1, \cdots, X_L) \in \mathbb{R}^{L\times d}$, $Y = (Y_1, \cdots, Y_L) \in \mathbb{R}^{L\times c}$ denote the sequence features of genomic loci in the context window and the corresponding RT signals, where $X_k \in \mathbb{R}^d$, $Y_k \in \mathbb{R}^c$ represent the feature vector and the RT signal of the $k$-th genomic locus within the context window, respectively. Here, $d$, $c$ denote the dimension of the feature vector for each locus and the number of types of genomic signals, respectively. Let $f_\theta : X \in \mathbb{R}^{L\times d} \to Y \in \mathbb{R}^{L\times c}$ be the function mapping features associated with a context window to the corresponding signals, with $\theta$ as the parameters. We have $c = 1$ in this work. The proposed model is applicable to data of multiple genomic signal types with $c > 1$. Suppose we select a subset of loci as the predictive loci from a context window. Let $\boldsymbol{\mathcal{V}}^{(L)} = \{\nu \in \mathbb{R}^L | \nu_k \in \{0, 1\}, k = 1, \cdots, L\}$. Let $S = (S_1, \cdots, S_L) \in \boldsymbol{\mathcal{V}}^{(L)}$ represent the selection of predictive genomic loci. $S$ is a $L$-dimensional binary value vector, where $S_k = 1$ if the $k$-th genomic locus is selected and $S_k = 0$ otherwise. Let $X_S$ be the representation of $X$ with only the features of selected genomic loci retained. Let $\odot$ represent element-wise multiplication. Suppose each element corresponds to a genomic locus. We have $X_S = X \odot S \in \mathbb{R}^{L\times d}$. Specifically, let $(X_S)_k$ denote the feature representation of the $k$-th locus in $X_S$. If $S_k = 0$, we set $(X_S)_k$ to be a zero-value feature vector. We have $(X_S)_k = \mathbf{0} \in \mathbb{R}^d$ if $S_k = 0$; $(X_S)_k = X_k$ if $S_k = 1$.

Given $X_S$, the mutual information between $X_S$ and $Y$ is:

$$I(X_S, Y) = \mathbb{E}\Big[\log \frac{p(X_S, Y)}{p(X_S)p(Y)}\Big] \tag{13}$$

$$= \mathbb{E}\Big[\log \frac{p(X_S)p(Y|X_S)}{p(X_S)p(Y)}\Big] = \mathbb{E}\Big[\log \frac{p(Y|X_S)}{p(Y)}\Big] \tag{14}$$

$$= \mathbb{E}\Big[\log p(Y|X_S)\Big] + \text{Constant} \tag{15}$$

Suppose $\phi_\beta : X \in \mathbb{R}^{L\times d} \to S \in \mathbb{R}^L$ is the function of the selector, mapping $X$ to $S$, with $\beta$ as the parameters. The objective function is:

$$\max_{\theta,\beta} \mathbb{E}\Big[\log p_\theta(Y|X_S)\Big], \tag{16}$$

$$\text{s.t. } S = \phi_\beta(X), \tag{17}$$

which can be rewritten as:

$$\max_{\theta,\beta} \mathbb{E}\Big[\log p_\theta(Y|X \odot \phi_\beta(X))\Big]. \tag{18}$$

Suppose the number of genomic loci is $N_g$ and the number of context windows is $N_c$. We assume that each genomic locus is the center of a context window. In this case, we have $N_g = N_c$. Specifically, suppose there are $N$ consecutive genomic loci without gaps, which are sequentially indexed with $1, \cdots, N$ in the order of their genomic coordinates. There are $N$ context windows centered on each genomic locus respectively, indexed with $1, \cdots, N$. Suppose the flanking region size on each side of a locus is $l$ genomic bins. For the $i$-th locus, the $(i-l)$-th - $(i+l)$-th loci are included in the corresponding $i$-th context window. We have $L = 2 \times l + 1$. Let $\boldsymbol{\mathcal{I}}_N = \{1, \cdots, N\}$. Let $x_i \in \mathbb{R}^{L\times d}$, $y_i \in \mathbb{R}^{L\times c}$ be the sequence features and signals of the $i$-th context window, respectively, $i \in \boldsymbol{\mathcal{I}}_N$. Let $\tilde{x}_k$, $\tilde{y}_k$ be the feature vector and the signal of the $k$-th genomic locus, respectively, $k \in \boldsymbol{\mathcal{I}}_N$. We have $x_i = (\tilde{x}_{i-l}, \cdots, \tilde{x}_i, \cdots, \tilde{x}_{i+l})$, $y_i = (\tilde{y}_{i-l}, \cdots, \tilde{y}_i, \cdots, \tilde{y}_{i+l})$. Here $x_i$ represents a context window consisting of the sequence features of $L$ associated genomic loci, while $\tilde{x}_k$ represents a single locus.

The context windows are overlapping. One genomic locus is involved in at most $L$ context windows, with the relative positions of the locus in the associated context windows ranging from 1 to $L$. Accordingly, in general the $i$-th locus can be in context with the $(i-2l)$-th locus to the $(i+2l)$-th locus. By using overlapping context windows, we involve the feature representation of a genomic locus in different contexts which are composed of varied combinations of genomic loci, extending the context scope of a single locus from $2l$ loci to at most $4l$ loci and utilizing more diverse context information to capture potential spatial dependencies across different subsets of loci.

The overlapping context windows can be formed by applying a sliding window along the genome with stride $d = 1$. If we use $1 < d < L$, we have $N_c < N_g$. In this case, not every genomic locus is the center of a context window, but every genomic locus is covered by at least one context window. Also, each context window is overlapped with at least one another context window. The learned model can still generate importance score estimates and RT signal predictions for each locus. In the prediction stage, we use the trained model to predict RT signals $\hat{y}_i \in \mathbb{R}^{L\times c}$ and estimate locus-wise importance scores $\alpha_i \in \mathbb{R}^L$ for the loci in each context window. For a locus that is involved in multiple context windows, we use the average of all the RT predictions for this locus in the associated context windows as the predicted RT signal, and we choose the importance score estimated in the context window where the locus is the center. In the case that $1 < d < L$ and $N_c < N_g$, we use the importance score averaged from the estimated scores for this locus in all the associated context windows.

##### *A.1.2 The selector and the predictor modules*

The proposed selector module performs the mapping $\phi_\beta : X \in \mathbb{R}^{L\times d} \to S \in \mathbb{R}^L$ with a neural network and a sampler which samples random variables from the Gumbel distribution. The selector module was described in **Methods** in the main text. Given the mapping function of the selector $\phi_\beta(X)$, of which the parameters $\beta$ are estimated and updated at each iteration, the objective function of the predictor module based on Eqn. 18 is:

$$\max_\theta \mathbb{E}\Big[\log p_\theta(Y|X \odot \phi_\beta(X))\Big], \tag{19}$$

where $p_\theta(Y|X)$ represents the probabilistic function of the predictor, with $\theta$ as the parameters. In the input layer of the predictor, the locus-wise sequence importance scores estimated by the selector are used to assign different weights to the sequence features of each genomic locus, which performs probabilistic approximation of binary sequence selection. The predictor aims to maximize the mutual information between the observed RT signals and the weighted sequence features in the context window. In the integrated framework of the two modules, we learn the parameters $\beta$ and $\theta$ simultaneously during the model training. The predictor module can be flexible and there can be different realizations of the module depending on the applications.

Our predictor module is constructed from four sequentially connected sub-modules. The first part is a preliminary feature transformation sub-module, with an interface from the selector module of the model.

The second part of the predictor module is a BiLSTM (bidirectional long short-term memory) layer [2], which is a variant of the LSTM model [3]). BiLSTM enables bidirectional sharing of context information across temporal or spatial series data. BiLSTM splits the hidden states of a standard LSTM into forward states and backward states, forming two parallel LSTM layers without interactions that enable the information flow in two directions. The output of BiLSTM is the concatenation of the forward hidden states and the backward hidden states. We utilize BiLSTM to model the bidirectional spatial dependencies across a sequence of genomic loci.

The third part is the self-attention layer as described in **Methods** in the main text. For a context window of size $L$, the estimated attention values can be represented by a matrix $A_L = \{a_{l,l'}\}_{l,l'\in\mathcal{C}_L} \in \mathbb{R}^{L\times L}$, where $a_{l,l'}$ represents the attention value locus $l'$ receives from locus $l$, measuring the importance of locus $l'$ in the prediction of the genomic signal for locus $l$. The fourth part is a fully-connected layer for RT signal prediction.

This method has the advantage that the number of model parameters does not increase with the context size, given the size of each genomic locus. Therefore, the method can be used to model spatial dependencies over very long distance without increasing model complexity.

##### *A.1.3 Model architecture and hyperparameters*

We implemented the model using Keras/Tensorflow. The architecture of the proposed model as implemented by Keras/Tensorflow is shown in **Fig.** S1. The main components of the selector module are as follows. For both the selector and the predictor modules, each of the dense layers and convolution layers is followed by a batch normalization layer [4] and an activation layer. The dense layer is applied to the locus-wise feature representations. The convolution layer is applied to the series of feature representations of the loci associated with the given context window.

1. A dense layer for preliminary feature transformation, followed by batch normalization and ReLU activation.
2. A 1D (one-dimensional) convolution layer to utilize local context information for importance estimation of each locus, followed by batch normalization and ReLU activation.
3. Dense layer for feature transformation, followed by batch normalization and ReLU activation.
4. Dense layer for estimating importance scores, followed by batch normalization and ReLU or sigmoid or linear activation.
5. A random variable sampler that samples random variables from Gumbel distribution Gumbel(0,1). Sampling are repeated $k$ times independently for each context window.
6. An element-wise Add layer to add the random variables from Gumbel(0,1) to the estimated importance scores. For each locus, there are $k$ values corresponding to the $k$ samplings. The maximal value of the $k$ results are retained for the corresponding locus.
7. Softmax layer following the element-wise Add layer to generate importance score estimates.

The hyperparameters of the different components we use in practice are as follows. Components (1,3,4): numbers of hidden units are 50, 25, 1, respectively. (2): number of filters: 50; kernel size: 3; stride: 1; padding method: 'same' (the left and right sides of the input are padded with zero values evenly such that the output has the same width dimension as the input). (5,6): $k = 10$, the temperature parameter $\tau = 0.1$.

The main components of the predictor module are as follows.

1. A dense layer for preliminary feature transformation, followed by batch normalization and ReLU activation.
2. A 1D convolution layer, followed by batch normalization and ReLU activation.
3. Dense layer for feature transformation, followed by batch normalization and ReLU activation.
4. An element-wise multiplication layer to multiply the feature representations of each locus with the importance scores estimated by the selector using the Gumbel-Softmax method.
5. BiLSTM layer. Layer normalization is applied.
6. Self-attention layer.
7. Dense layer, followed by batch normalization and sigmoid or tanh activation to predict the genomic signals.

The hyperparameters of the different components we use in practice are as follows. Components (1,3,7): numbers of hidden units are 50, 50, 1, respectively. (2): number of filters: 50; kernel size: 3; stride: 1; padding method: 'same'. (5): number of hidden units in each single LSTM layer: 32; activation function: tanh. (6): number of hidden units: 50; activation function: tanh.

We used the Adam optimizer. The batch size is 512. The learning rate is 0.0005. We chose the $l_1$ regularization coefficients to be zero. We chose the $l_2$ regularization coefficients to be 1e-04 or 1e-05. The regularization is applied to both the selector and predictor modules. The parameter configurations were selected based on searching through a range of combinations of different parameters with evaluation on the validation data. We used early stopping where training is stopped if no performance gain is observed on the validation data for consecutive 7 epochs.

For hyperparameter configuration of CONCERT, we used grid-search over specific ranges of specific hyperparameters. Specifically, we tuned the number of kernels and the kernel size in each convolution layer, the number of hidden units in each dense layer, the number of times to perform Gumbel-Softmax based predictive loci sampling by the selector (the parameter $k$), the temperature parameter $\tau$ used for the Gumbel-Softmax trick, the number of hidden units in the BiLSTM layer of the predictor, the loss function of the predictor, and the learning rate in CONCERT. We selected the hyperparameter configurations based on performance evaluation on the validation data.

###### *A.1.4 Feature representation of the genomic loci*

With prior knowledge of the possible dependencies between DNA sequence and RT signals, we generate different types of features for each genomic bin. We concatenate different types of features to form a feature vector of the corresponding genomic bin, which is fed to the model as input. The candidate feature types include the $K$-mer frequency based features, the GC profile based features, the TF binding motif frequency based features (optional), and the evolutionary information based features (optional). Let $x_i^{(\text{kmer})}$, $x_i^{(\text{gc})}$, $x_i^{(\text{motif})}$, and $x_i^{(\text{phyloP})}$ be the corresponding types of features of the $i$-th genomic locus, respectively, $i = 1, \cdots, N$. The performance evaluation of RT prediction in the cell line H1-hESC using different combinations of the different types of feature representations are shown in **Table** S5. In this work, we primarily used the $K$-mer frequency based features and the GC profile based features, as the combination of the two feature types show higher prediction performance (**Table** S5) and they are also simpler in computation. The concatenated feature vector of genomic locus $i$ is $x_i = (x_i^{(\text{kmer})}, x_i^{(\text{gc})})$.

(1) *Feature representation based on $K$-mer frequency.* We extract $K$-mer based features from the DNA sequence of each genomic locus. Given the four types of nucleotides: $A$, $G$, $C$, $T$, there are $4^K$ different types of $K$-mers for a specific $K$. For a given sequence of a genomic locus, the number of occurrences of each type of $K$-mer, normalized by the sequence length, forms a $4^K$-dimensional feature vector. We choose $K = 5$ and $K = 6$, and concatenat the corresponding feature vectors $\tilde{x}_i^{(\text{kmer},5)}$ and $\tilde{x}_i^{(\text{kmer},6)}$ for the $i$-th locus. We have $\tilde{x}_i^{(\text{kmer})} = (\tilde{x}_i^{(\text{kmer},5)}, \tilde{x}_i^{(\text{kmerr},6)}) \in \mathbb{R}^{5120}$. The $K$-mer frequency based feature vector is high-dimensional. We evaluated different feature dimension reduction methods based on the prediction performance on the validation data using the corresponding transformed features. The methods we compared include PCA, Kernel PCA [5], Sparse PCA [6], ICA (Independent Component Analysis) [7], Mini-Batch Dictionary Learning [8], Truncated SVD (singular value decomposition) [9], and Autoencoder [10]. We use the scikit-learn library [11] for implementation of the methods.

We pooled the training and test samples (each sample is a genomic locus), and performed feature dimension reduction on the pooled data using each of the methods above, without using RT signal information of the samples. We compared the RT prediction performance using features from the different corresponding methods. Based on the evaluation of the prediction performance and the computation efficiency, we choose PCA for the dimension reduction of the $K$-mer frequency features and choose the reduced dimension to be $d_k = 50$. For the PCA method, we use the top $d_k$ components with the largest variances on the transformed coordinates in the feature space after dimension reduction as the feature values of the corresponding genomic locus.

(2) *Feature representation using GC profile.* The GC content of DNA sequences is known to correlate with RT profiles [12]. Early replications were found to be enriched in GC-rich regions. We extracted two types of GC-based features from the sequence of each genomic locus, which are GC content (denoted by $f^{(\text{gc})}$) and GC skew (denoted by $f^{(\text{skew})}$), respectively. Let $n(\alpha)$ be the number of nucleotide $\alpha$ in a given

DNA sequence, $\alpha \in \{A, G, C, T\}$. For a given genomic locus, we have

$$f_i^{(\text{gc})} = \frac{n(G) + n(C)}{n(A) + n(G) + n(C) + n(T)}, \tag{20}$$

$$f_i^{(\text{skew})} = \frac{n(G) - n(C)}{n(G) + n(C)}. \tag{21}$$

$f^{(\text{gc})} \in [0, 1]$, $f^{(\text{skew})} \in [-1, 1]$. Let $x_i^{(\text{gc})}$, $f_i^{(\text{gc})}$, $f_i^{(\text{skew})}$ be the GC-based feature vector, the GC content feature, and GC skew feature of the $i$-th genomic locus, respectively. We have $x_i^{(\text{GC})} = (f_i^{(\text{gc})}, f_i^{(\text{skew})})$.

(3) *Feature representation using transcription factor binding motifs.* We use TF binding motif frequency as an optional feature type for the genomic loci. We used the software FIMO [13] and 769 position weight matrices (PWMs) of TF binding motifs from the HOCOMOCO v11 database [14] to perform motif scanning on human genome hg38. In each locus, we counted the motif frequency for each PWM ($P$<1e-05 required for each motif) and normalized the frequency by the genomic locus size, obtaining a 769-dimensional feature vector. We then performed dimension reduction on the motif frequency features. Since the motif frequency features are sparse and the truncated SVD method [9, 15] is appropriate for transformation of sparse features, we used truncated SVD for dimension reduction of the motif features, with the new feature dimension to be $d_m = 50$.

(4) *Feature representation using phyloP scores.* We downloaded the phyloP (phylogenetic $p$-values) scores on the human genome from the UCSC Genome Browser. The phyloP scores are conservation scores calculated by phyloP [16] from the PHAST package [17] based on multiple sequence alignments of 99 vertebrate genomes to the human genome hg38, denoted as 100-way phyloP scores. A phyloP score is assigned per base pair in most regions of the human genome. For each genomic locus, we extracted a feature vector from the phyloP scores in this locus by calculating different types of statistics of the scores. Suppose $c_i = \{c_{i,1}, \cdots, c_{i,N_i}\}$ are the phyloP scores in the $i$-th genomic locus, where $N_i$ is the number of phyloP scores in this locus. First, we calculated the average, median, maximum and minimum of the scores in $c_i$. Second, we calculated the distribution of the scores across different intervals. Given that the observed phyloP scores ranging from -20 to 10, we divide the range [-20,10] into 15 evenly spaced non-overlapping intervals with length of 2 each. We calculated the frequency of phyloP scores in each interval, normalized by $N_i$. Let $\{x_{i,k}^{(\text{phyloP})}\}_{k=1}^{4}$, $\{x_{i,k}^{(\text{phyloP})}\}_{k=5}^{19}$ be the four statistics from the first step and the 15 normalized frequencies from the second step, respectively. Let $x_i^{(\text{phyloP})}$ be the feature vector based on the phyloP scores for the $i$-th locus. We have $x_i^{(\text{phyloP})} = (x_{i,1}^{(\text{phyloP})}, \cdots, x_{i,19}^{(\text{phyloP})})$.

##### *A.1.5 Hierarchical structure to learn feature representation from genomic sequences*

We also developed a hierarchical structure of CONCERT (denoted as CONCERT-hierarchical) as an alternative to the basic CONCERT model to learn feature representation from locus-wise genomic sequences. The architecture of CONCERT-hierarchical consists of two levels. The first level is a CNN-BiLSTM sub-model applied to each locus in the context window, to perform local feature representation learning of locus-wise sequences, by learning spatial dependencies across nucleotides in the sequence of each locus. The second level is the same as the basic CONCERT model without the input layer for pre-engineered feature representation, consisting of the selector module and the predictor module. The input to the first level is one-hot encoded feature vectors of the sequences from each locus in each context window. The output from the first level is fed to the second level, to replace the original pre-engineered input layer.

Suppose there are $L$ genomic loci in each context window, with the genomic bin size of $w$. The input feature vector of each locus is a one-hot encoded sequence, which is a binary-valued matrix of dimensions $4 \times w$, where each row corresponds to the nucleotide type $\alpha \in \{A, C, G, T\}$ and each column represents a base in the sequence. For each column, the row corresponding to the nucleotide at this position is set to 1, and the other rows are zeros. The input feature of each context window is $(4 \times w \times L)$-dimensional. The structure of the CNN-BiLSTM sub-model is as follows:

1. Three sequentially connected convolution layers, each followed by batch normalization, ReLU activation, and a max-pooling layer. The number of kernels in each convolution layer is 128, 32, 32, respectively. The kernel size of each layer is 10, 7, 5. The stride of convolution operations in each layer is 5, 1, 1. The padding method for each layer is "valid" (input is not padded on each side). The size of max pooling following each convolution layer is 10, 5, 2, with the strides 10, 5, 2, respectively.
2. One BiLSTM layer. The number of hidden units in each single LSTM layer of the BiLSTM layer is 32. Layer normalization is applied. The activation function is tanh.
3. A dense layer, followed by batch normalization and ReLU activation. Number of hidden units: $50$.
4. A global max-pooling layer or a concatenation layer. If a global max pooling layer is used, max pooling is applied to the outputs of the dense layer for the loci in a context window to form a $d$-dimensional feature vector of the window. If concatenation is used, the outputs of the dense layer for each locus in the context window are concatenated to form a feature vector of dimension $w \times d$.

We use CONCERT-basic to denote the CONCERT model using pre-engineered feature representations of genomic loci. The RT prediction performance of CONCERT-hierarchical is similar to but lower than the performance of CONCERT-basic in H1-hESC cell line. Also, the time and space consumption costs of CONCERT-hierarchical are higher than those of CONCERT-basic. We found that feature learning from local genomic sequences did not show an advantage over using the pre-engineered features in our study (**Table** S6). A possible reason is that the combination of feature learning for genomic bins of 5Kb in size and modeling spatial dependencies across genomic bins over long distance increases the model complexity. The model is more likely to overfit and get trapped in local minima. We therefore primarily use the prediction results of CONCERT-basic for subsequent analysis.

#### A.2 Performance evaluation of CONCERT in comparison with other methods

##### *A.2.1 Model training and prediction in mESCs and human cell lines*

We performed cross-chromosome model training and prediction. We used two non-overlapping subsets of chromosomes for training. For each subset of chromosomes, the other chromosomes not included in the subset were used as chromosomes for testing. We used the chromosomes in each subset for training and made predictions with the trained model on the left-out chromosomes. Specifically, for each cell type in human, we used chromosome 1 to chromosome 16 as alternating training and test data. We used chromosome 17 to chromosome 22 as reserved test data. We only included autosomes in this work. We chose the context of +/-50 bins (bin size: 5Kb) for each genomic locus (genomic bin) to form a context window centered at this locus. Therefore, the context size of each locus is 505Kb (101 bins). Specifically, for each cell type, first we used the odd-numbered chromosomes of chromosome 1 (with the abbreviated notation chr1) to chromosome 15 (chr1, chr3, $\cdots$, chr15) for model training, and predicted RT signals on the left-out 14 autosomes with the trained model. Next, we used the even-numbered chromosomes of chr2 to chr16 (chr2, chr4, $\cdots$, chr 16) for training, and predicted RT signals on the left-out autosomes.

We split the samples on the chromosomes used for training into training data and validation data, with the ratio 9:1. Specifically, for each chromosome in the training set, we selected the first 90% of genomic bins for training based on the genomic coordinates, and used the last 10% of genomic bins as validation data. With the described training and prediction scheme, we obtained RT signal predictions on all the autosomes. For RT prediction in mESCs, for each of the two alleles (mutant and WT), we first used the odd-numbered chromosomes of chr1 to chr17 as the training data. We used the trained model to predict RT signals and estimate importance scores on the left-out 11 autosomes, including chromosome 8 and chromosome 16 which differ in the presence of the identified ERCEs between the two alleles. Next, we used the even-numbered chromosomes of chr2 to chr18 for model training, and performed RT prediction and importance score estimation on the left-out autosomes.

For model training and prediction, the RT signals were scaled to [0,1]. First, we processed the raw Repli-seq data to retrieve RT signals at each genomic locus as described in **Data collection and processing** in **Methods**. For signal normalization, we set a upper bound and a lower bound of the RT signals and truncated the RT signals between the bounds, in order to reduce the influence of extreme high or low signal values on the normalization. Specifically, we identified the highest and lowest RT signals on each chromosome. We then used the median of the highest or lowest RT signals across the chromosomes as the upper bound or the lower bound, respectively. The RT signals above the upper bound or below the lower bound were set to the upper or lower bound, respectively. For RT signals in the training data, only chromosomes that were used for training were involved in estimating the upper and lower bounds. The RT signals were then normalized to the scale [0,1] by minmax scaling, with the minimal value corresponding to 0 and the maximal value corresponding to 1.

##### *A.2.2 RT prediction performance evaluation*

The evaluation metrics we used for comparing the predicted RT signals with the ground truth RT signals include Pearson correlation coefficient (PCC, or Pearson's $r$), Spearman's rank correlation coefficient (Spearman's $\rho$), the explained variance, and the coefficient of determination ($R^2$ score).

Let $\hat{Y}$, $Y$ denote the predicted RT signals and the ground truth RT signals, respectively. For predictions on a sample of $N$ genomic loci, let $\hat{y}$, $y$ denote the predicted signals and the ground truth signals, respectively. We have $\hat{y} = \{\hat{y}_1, \cdots, \hat{y}_N\}$, $y = \{y_1, \cdots, y_N\}$.

The Pearson correlation coefficient is defined as:

$$\rho_{Y,\hat{Y}} = \frac{\mathbb{E}(Y\hat{Y}) - \mathbb{E}(Y)\mathbb{E}(\hat{Y})}{\sqrt{\mathrm{Var}(Y)}\sqrt{\mathrm{Var}(\hat{Y})}}, \tag{22}$$

where $\mathrm{Var}$ represents the variance of a random variable.

The Spearman's rank correlation coefficient (denoted as $r_s$) is defined as the Pearson correlation coefficient between two rank variables. Let $rg_Y$, $rg_{\hat{Y}}$ be the rank variables converted from $Y$, $\hat{Y}$, respectively. For a sample $y = \{y_1, \cdots, y_N\}$, we have $rg_Y = \{rg_{y_1}, \cdots, rg_{y_i}\}$, where $rg_{y_i}$ is an integer representing the rank of $y_i$ in the set of $\{y_1, \cdots, y_N\}$ based on the predefined order of the values, $i = 1, \cdots, N$. The same definition applies to $rg_{\hat{y}_i}$. The Spearman's rank correlation coefficient is defined as:

$$r_s(Y, \hat{Y}) = \rho_{rg_Y, rg_{\hat{Y}}}, \tag{23}$$

where $\rho$ represents the Pearson correlation coefficient between two random variables.

The Pearson correlation coefficient and the Spearman's rank correlation coefficient are both in the range of [-1,1], with the higher value representing better prediction performance.

The explained variance is defined as:

$$\text{explained\_variance}(Y, \hat{Y}) = 1 - \frac{\text{Var}(Y - \hat{Y})}{\text{Var}(Y)}, \tag{24}$$

The $R^2$ score is estimated as:

$$R^2(y, \hat{y}) = 1 - \frac{\sum_i^N (y_i - \hat{y}_i)^2}{\sum_i^N (y_i - \bar{y})^2}, \tag{25}$$

where $\bar{y} = \frac{1}{N} \sum_i^N y_i$. Both of the explained variance and the $R^2$ score are in the range of $(-\infty, 1]$, with the higher value representing the better prediction performance.

For RT classification, genomic loci with original RT signals above the threshold zero is labeled with 1 (early replication) and otherwise labeled with 0 (late replication). For evaluation of early/late RT classification performance, we used the metrics AUROC (area under receiver operating characteristic curve) and AUPR (area under precision-recall curve). We normalized the predicted RT signals to the scale of [0,1], with the higher value corresponding to earlier replication.

##### *A.2.3 Predicting RT signals using sequence features without context*

We evaluated the prediction performance of our method in comparison with six methods. Five of the methods extract features from the sequence of each locus without utilizing the context information. The first method is XGBoost regression (XGBR). XGBboost regression utilizes Gradient Tree Boosting to fit regression models for continuous value variables. The second method is random forest (RF). The third method is linear regression (LR). The fourth method is a locus-wise deep neural network model (noted as DNN-local). DNN-local is similar to the predictor module of CONCERT, except that the convolution layer is replaced by a dense layer (or a convolution layer with kernel size of 1), the BiLSTM layer is replaced by two connected dense layers each with the same number of hidden units as the single LSTM layer of the BiLSTM, and there is no self-attention layer. The input to each of the four methods are pre-engineered feature representations of each genomic locus based on transformed *K*-mer frequency and GC-content features, as described in **Supplementary Methods** A.1.4.

The fifth method is adapted from the model of DanQ [18]. The model consists of three sequentially connected components, including multiple convolution layers to extract intermediate features from the sequence of each locus, a BiLSTM layer to capture spatial dependencies across nucleotides in the sequence of the corresponding locus for higher-level feature representation, and a dense layer to predict the corresponding RT signal. Different from the other methods, the DanQ-adapted model does not utilize pre-engineered feature representations, but instead performs genomic signal prediction *de novo* from the sequence of each locus. We name the fifth method as CNN-BiLSTM-local.

The five methods use feature representation from the sequence of each individual locus to predict the corresponding genomic signal, without modeling spatial dependencies across genomic loci in the larger-scale genomic regions.

Specifically, the model structure of DNN-local is as follows. (1) Three sequentially connected dense layers, each followed by batch normalization and ReLU activation. Number of hidden units per layer: 50. (2) Two dense layers, each followed by batch normalization and ReLU activation. Number of hidden units per layer: 32. (3) A dense layer, followed by batch normalization and sigmoid activation. Number of hidden units: 1.

The model structure of CNN-BiLSTM-Local is as follows. (1) Three sequentially connected convolution layers, each followed by batch normalization, ReLU activation, and a max-pooling layer. (2) A BiLSTM layer. (3) A dense layer (optional), followed by batch normalization and ReLU activation. (4) Self-attention layer (optional). (5) A global max-pooling layer or a concatenation layer to transform the 2D output of the previous layer into a 1D feature vector. (6) A dense layer, followed by batch normalization and ReLU activation. (7) A dense layer, followed by batch normalization and sigmoid activation. The hyperparameters of the different components we use in practice are as follows. (1): the number of kernels in each convolution layer is 10, with the kernel size 5; the max pooling size is 5, with the stride 5. (2): number of hidden units in each single LSTM layer is 32. (3,6,7): number of hidden units are 50, 50, 1, respectively. (4): number of hidden units is 50; activation function is tanh.

We performed hyperparameter tuning for the compared methods using grid-search over specific ranges of hyperparameters, based on prediction performance on the validation and test data. For XGBoost and RF, we tuned the number of trees, the max depth of each tree, and the learning rate each within a specific range. For DNN-local, the model structure partially reassembles that of the predictor module of CONCERT, with the difference that adjacent genomic loci are disconnected in the modeling. Accordingly, the hyperparameters in the predictor module of CONCERT are used with adjustment in DNN-local, as described above. For CNN-BiLSTM-local, we tuned the number of kernels and the kernel size in each convolution layer, the max pooling sizes, and the number of hidden units in the BiLSTM layer. We also compared the performance between using global max-pooling and using concatenation to generate locus-wise feature vectors from the output of the BiLSTM layer. The performances are similar for the different configuration variants we tested, which did not show change of the relative performance disadvantage compared to the methods using the context information of sequences.

##### A.2.4 Predicting RT signals using sequence features with dilated convolutions

The sixth method for RT prediction performance comparison is named Dilated-CNN, which is adapted from the module of densely connected dilated convolution layers in the model of Basenji [19]. Dilated-CNN shares the identical input with CONCERT, using overlapping context windows with pre-engineered feature representations of genomic loci within each window.

The Basenji model consists of two main modules. The first module utilizes sequential convolution layers and max pooling layers to generate one feature vector for each genomic bin from the corresponding sequence. The second module takes advantage of multiple densely connected dilated convolution layers to capture the spatial dependencies across genomic bins in a long sequence. Dense connection represents that each dilated convolution layer takes the output of all the previous layers in the module as input. The gap size of the dilated convolution kernel increases exponentially across the sequential dilated convolution layers. Dilated-CNN reassembles the second module of Basenji. In Dilated-CNN, we assume that the feature representation of each genomic locus has been generated through feature engineering, and we employ the same structure of the densely connected dilated convolution layers in Basenji to model long-range spatial dependencies across loci within a context window. Specifically, we use seven densely connected dilated convolution layers, with the dilation rate (measuring the gap size of a kernel) increasing exponentially from 1 to 64, and the kernel size of each layer as 3.

For Dilated-CNN, we turned the number of densely connected dilated convolution layers and the kernel size per layer, with number of layers ranging from 4,5,6,7 and kernel size changing between 3 and 5. The original parameter configuration used in Basenji (7 dilated convolution layers with kernel size of 3 per layer) reaches relatively higher prediction accuracy measured by Pearson correlation coefficient on average across the different cell lines. We use the original parameter configuration accordingly.

##### A.2.5 Performance comparison of CONCERT and dilated convolution-based models

The performance comparison between CONCERT-basic and Dilated-CNN is shown in **Fig.** S2, **Fig.** S3, and **Table** S2. For performance comparison of CONCERT-hierarchical with dilated convolution-based models, we adapted the Basenji model [19] to perform RT signal prediction from genomic sequences without using pre-engineered feature representations. The original Basenji model was designed for regulatory activity prediction from genomic sequences, using the genomic bin size of 128bp, which is not directly applicable to RT prediction (genomic bin size of 5Kb) in our study. To have comparable prediction results, we tailed the Basenji model towards RT prediction. Specifically, we use four sequentially connected convolution layers each followed by a max-pooling layer to generate a feature vector for each genomic bin within each context window. The features of loci in each context window are fed to seven densely connected dilated convolution layers, using the same configurations as used in Basenji. We denote the model as Basenji-adapted.

Specifically, for the CNN module of Basenji-adapted, the number of kernels in each convolution layer in the CNN module of Basenji-adapted is 128, 32, 32, 32, respectively. The kernel size of each layer is 10, 7, 5, 5. The stride of convolution operations in each layer is 5, 1, 1, 1. The padding method for each layer is “valid”. The size of max pooling following each convolution layer is 10, 5, 2, 2, with the strides 10, 5, 2, 2, respectively.

CONCERT-hierarchical and Basenji-adapted both predict RT signals *de novo* from base-pair resolution genomic sequences utilizing long-range context information, without pre-engineered features (**Supplementary Methods** A.1.5). We compared the RT prediction performances of Basenji-adapted and CONCERT-hierarchical in H1-hESC cell line. The Pearson correlation coefficient (PCC), Spearman’s rank correlation coefficient (Spearman’s $\rho$) between the real RT signal and the predicted RT signal by Basenji-adapted are 0.65,0.72, respectively. The PCC and Spearman’s $\rho$ between the real RT signal and the predicted RT signal by CONCERT-hierarchical are 0.79, 0.79, respectively. Basenji-adapted has lower performance than Dilated-CNN, which is adapted from the dilated convolution module of Basenji [19]. Also, the time and space costs of both CONCERT-hierarchical and Basenji-adapted are higher than the basic CONCERT model which uses pre-engineered feature representations of genomic loci.

#### A.3 Comparison between the CONCERT predicted important loci and other genomic and epigenomic features

##### A.3.1 Processing importance scores for RT-modulating genomic loci identification

We generated a set of annotations of the potentially predictive genomic loci based on the locus-wise importance scores estimated by our method on the Repli-Seq data of each cell type. We normalized the estimated importance scores on each chromosome to the scale of [0,1], based on the rank of the score of each locus among all the loci on the corresponding chromosome. The original score not below $x\%$ of all the scores is transformed to be $x \in [0, 1]$. We then filtered the genomic loci to select a subset of loci as potentially predictive genomic loci. First, we performed peak calling on the importance scores along each chromosome using the peak finding function in the scikit-learn library [11], with constraints on both the peak strength and the minimal distance between two adjacent peaks, to suppress local non-maximal scores. Specifically , we require the normalized estimated score of a peak to be above 0.90. Let $S_1$ denote the set of loci with identified local peaks of the estimated scores. Second, we select genomic loci with the normalized score above 0.975, denoted as set $S_2$. The union of $S_1$ and $S_2$ is the selected subset of predictive genomic loci. We divide the selected loci into early RT group (denoted by $S_E$) and late RT group (denoted by $S_L$) based on the early or late RT domains they reside in, respectively. In this study we mainly analyze loci in $S_E$.

##### A.3.2 CONCERT *estimated importance score comparison between ERCEs and non-ERCEs*

We compared the estimated importance scores between the ERCEs and the non-ERCE genomic regions in the mESC CAST/129 hybrid cells. We only included the predicted ERCEs on autosomes in the analysis. For each ERCE, we randomly sample 200 genomic regions of the same length as the ERCE and without overlapping with any ERCE from the genome. An ERCE ranges from 50Kb to 200Kb in length, containing 10-40 genomic bins. For each ERCE or each randomly sampled element, we calculated both the maximal value and the mean value of the estimated importance scores of genomic bins in the ERCE or the sampled region. We then merged the maximal values (or mean values) of the estimated importance scores associated with each randomly sampled region to form the background importance score distribution. We compared the distribution of the estimated maximal-value (or mean-value) importance scores of ERCE in comparison with the background distribution. ERCEs have higher maximal-value (or mean-value) per-region importance scores than the randomly sampled genomic regions. To map mESC ERCEs to the human genome, we used liftOver [20] with the minimal remapping ratio of 0.5 (mapping between mm10 and hg38), requiring the regions mapped to the human genome can be mapped back to the mouse genome and overlap with the original predicted ERCE regions (i.e., reciprocal liftOver). To identify mapped ERCEs with pluripotency TF co-binding, we used the binding site annotations of POU5F1, SOX2, and NANOG based on ChIP-seq data from the ENCODE project [21] and [22].

##### A.3.3 *Comparing the predicted important genomic loci with cis-regulatory elements*

To evaluate the estimated importance scores of genomic loci with *cis*-regulatory elements (CREs) in open chromatin regions, we downloaded peak region annotations of DNase-seq data in each of the five cell lines from the ENCODE project [21]. We extended each peak by +/-250bp. Among all the genomic loci that are overlapping with the extended DNase-seq peak regions (noted as open chromatin regions), we compared the distributions of the estimated importance scores between the loci that contain at least one CRE and the loci that do not overlap with any CRE. We used the importance score distribution of the non-CRE genomic loci as the background distribution.

To calculate the enrichment of TF binding site in the predicted important genomic loci, we used the peak regions identified from the ChIP-seq data of the corresponding TFs from the ENCODE project [21]. We sampled background loci for the predicted important genomic loci with matched RT signal levels (defined in **Supplementary Methods** A.3.4). Let $N_p$, $N_b$ be the number of predicted important loci and the number of background loci in the open chromatin regions, respectively. Let $N_p^{(k)}$, $N_b^{(k)}$ be the number of predicted important loci and the number of background loci that overlap with the binding site of protein $k$ in the open chromatin regions, respectively. We performed Fisher's exact test based on the contingency table formed by $\{N_p, N_b, N_p^{(k)}, N_b^{(k)}\}$ to evaluate if the binding site of TF $k$ is significantly enriched in the predicted important loci in the open chromatin regions, using the threshold of $P<0.05$.

##### A.3.4 *Sampling background genomic loci for the predicted important loci*

To evaluate the enrichment of specific sequence features in the predicted important loci, we sampled background loci which match the predicted loci in terms of RT signal levels. Specifically, the genomic loci were assigned to four groups based on the the corresponding RT signal values. The loci with RT signals ranking top 50% among all the loci in the early RT regions were assigned to group 1, and the other early RT loci were assigned to group 2. We sorted the loci in late RT regions by the corresponding RT signals in ascending order. The loci ranking top 50% in the late RT regions were assigned to group 4, and the other late RT loci were assigned to group 3. Group 1 and group 4 correspond to the loci with strong early RT and strong late RT signals, respectively. We then randomly sampled background loci for each predicted important locus within the same group. Therefore, the sampled background loci matched the predicted important locus in terms of RT signal levels.

##### A.3.5 Chromatin state annotations by ChromHMM

To analyze the enrichment of the different chromatin states in the predicted important genomic loci for RT and to identify the genomic loci without known regulatory elements, we used the chromatin states estimated by ChromHMM [23]. There are 15 ChromHMM states: TssA (Active Transcription start site, Active TSS), TssAFlnk (Flanking Active TSS), TxFlnk (Transcription at gene $5'$ end and $3'$ end, TX (Strong transcription), TxWk (Weak transcription), EnhG (Genic enhancers), Enh (Enhancers), ZNF/Rpts (ZNF genes and repeats), Het (Heterochromatin), TssBiv (Bivalent/Poised TSS), BivFlnk (Flanking Bivalent TSS/Enh), EnhBiv (Bivalent Enhancer), ReprPC (Repressed PolyComb), ReprPCWk (Weak Repressed PolyComb), and Quies (Quiescent/Low). We further merged the 15 ChromHMM states into 8 groups, following the definition in [24]. Specifically, the 8 groups are (i) BIV: Bivalent, including the states TssBiv and BivFlank; (ii) ENH: Enhancers, including EnhBiv, EnhG, and Enh; (iii) TSS: Transcription start site, including TssA and TssAFlnk; (iv) TX: Transcription, including TxFlnk, Tx, and TxWK; (v) REPEAT: Repeats, including ZNF/Rpts; (vi) REPRESS: Repressed, including ReprPC and ReprPCWk; (vii) HET: Heterochromatin, including Het; (viii) QUIES: Quiescent, including Quies.

##### A.3.6 Repetitive element enrichment analysis of the predicted important loci

We compared the predicted important loci with transposable elements (TEs) and repetitive element (RE) (**Figs.** S7-S8). Using the human genome repetitive sequence annotations retrieved from the UCSC Genome Browser [25], we calculated the enrichment of each TE/RE at different importance score levels. Specifically, For each human cell type, we classified the genomic loci into $K = 20$ groups of equal sizes based on the ranking of the estimated importance scores of the loci in the descending order. Each group corresponds to an importance score level. We then calculated the fold change of the coverage of each TE/RE family in each importance group in comparison with the average coverage across all groups.

##### A.3.7 Identifying genomic loci depleted of known regulatory elements

To identify the genomic loci without regulatory element annotations in the studied human cell lines, we used the following annotation data: (i) ChromHMM state annotation [23]. There are 15 ChromHMM states, the notations of each state have been described in **Supplementary Methods** A.3.5. If a predicted important genomic locus (with +/-1 bin (5Kb) extension) overlaps with any of the following ChromHMM states that are directly related to transcriptional activities or regulatory elements, we consider it as possibly containing regulatory elements. The corresponding specific ChromHMM states are: TssA, TssAFlnk, TssBiv, TxFlnk, TX, TxWk, BivFlnk, EnhBiv, EnhG, and Enh. (ii) candidate CRE annotation retrieved from the SCREEN database [26]. There are five main CREs, including dELS (distal enhancer-like signatures), pELS (proximal ELS), PLS (promoter-like signatures), DNase-H3K4me3 (the likely poised elements with DNase and H3K4me3 marks), and CTCF-only, respectively. If a predicted important genomic locus overlaps with any of the five types of CREs, we consider it as possibly containing regulatory elements. (iii) Enhancer annotation retrieved from the EnhancerAtlas 2.0 databases [27].

We retain the predicted important loci that do not overlap with an annotated enhancer in the corresponding cell line. The original enhancer annotations were based on the hg19 genome and we mapped the annotations to the hg38 genome using liftOver. If a predicted important locus (with +/-1 bin extension) is not overlapped by any of the annotations from (i,ii,iii), we consider the locus as without known regulatory elements.

###### A.3.8 Gene expression analysis for the predicted important genomic loci

We compared the potentially associated gene expression levels between the predicted RT-predictive important genomic loci and the background loci in cell lines H1-hESC, GM12878, and K562, using the gene expression data downloaded from the ENCODE project [21]. First, we identified the genes with cell type-specific expression or cell type-specific increased expression. Specifically, for each of the compared cell types H1-hESC, GM12878, and K562, if a gene has TPM (transcript per million)$>1$ in the corresponding cell type and TPM$<1$ in the other cell types, we consider it as a cell type-specific expressed gene. For the remaining genes with TPM$>1$ in this cell type, if the gene expression level measured by TPM is at least 2 times the gene expression level in each of the other two cell types, we consider it as a gene with cell type-specific increased expression. We then identified cell type-specific expressed genes in proximity to the predicted important loci, including cell type-specific important loci. If a gene is within specific distance (+/-5Kb) to a predicted important locus, we consider the gene as in proximity to the locus and possibly associated with the locus.

###### A.3.9 Hi-C data analysis

We performed Hi-C aggregate analysis of the predicted important loci in different human cell types, including H1-hESC, HCT116, IMR90, and K562. The Hi-C data were downloaded from 4DN data portal [28]. To measure the aggregate enrichment of predicted important loci in the Hi-C contact matrix, we calculated the sum of intra-chromosomal Hi-C contacts over submatrices centered at paired predicted important loci interactions. The size of the submatrix is 21 x 21 square at the resolution of 25kb (each block in the submatrix represents a 25kb bin), and the centers of the submatrices correspond to off-diagonal coordinates for the corresponding pairs of predicted important loci. For comparison, we performed the same Hi-C aggregate analysis for the background loci (the control group), which were sampled to match the predicted important loci in RT signal levels (defined in **Supplementary Methods** A.3.4).

We analyzed the binding score distribution of CTCF in genomic regions harboring the predicted important loci in different human cell types, using the CTCF ChIP-seq data downloaded from ENCODE. The fold change score from the ChIP-seq data corresponds to the strength of a binding event of the CTCF protein. We calculated the average of CTCF binding scores across the genomic loci as a function of distance to a predicted important locus within the range of +/-100kb.

We used the HiCCUPS program from the Juicer tool [29] to detect Hi-C loops. For H1-hESC cell line, where multiple protocol variations of *in situ* Hi-C data are available, we used data using restriction enzyme DpnII, without and with FA&DSG treatment (referred to as Hi-C 1.0 and Hi-C 2.0). In addition to Hi-C, we also obtained Micro-C data for H1-hESC cell line and identified Micro-C loops as additional evidence for chromatin interactions [30]. All loops were detected at 10kb or 25kb resolution with default parameters in HiCCUPS program.

We calculated the Hi-C O/E (observed/expected) interaction score for each pair of predicted important genomic locus and a RT origin identified from SNS-seq data within a distance range. For a pair of predicted important locus and RT origin, we randomly sampled a locus from the set of background loci (defined in **Supplementary Methods** A.3.4) to pair with the RT origin, with the distance between the locus and the RT origin following the same distribution of the distance between the RT origin and the predicted important locus to calculate the expected Hi-C O/E scores, used as the control group.

[21] ENCODE Project Consortium. An integrated encyclopedia of DNA elements in the human genome.

Nature. 2012;489(7414):57-74.
[22] Lister R, Pelizzola M, Dowen RH, Hawkins RD, Hon G, Tonti-Filippini J, et al. Human DNA methylomes at base resolution show widespread epigenomic differences. Nature. 2009;462(7271):315-22.
[23] Ernst J, Kellis M. ChromHMM: automating chromatin-state discovery and characterization. Nature Methods. 2012;9(3):215-6.
[24] Grubert F, Srivas R, Spacek DV, Kasowski M, Ruiz-Velasco M, Sinnott-Armstrong N, et al. Landscape of cohesin-mediated chromatin loops in the human genome. Nature. 2020;583(7818):737-43.
[25] Casper J, Zweig AS, Villarreal C, Tyner C, Speir ML, Rosenbloom KR, et al. The UCSC Genome Browser database: 2018 update. Nucleic Acids Research. 2017;46(D1):D762-9.
[26] Moore JE, Purcaro MJ, Pratt HE, Epstein CB, Shoresh N, Adrian J, et al. Expanded encyclopaedias of DNA elements in the human and mouse genomes. Nature. 2020;583(7818):699-710.
[27] Gao T, Qian J. EnhancerAtlas 2.0: an updated resource with enhancer annotation in 586 tissue/cell types across nine species. Nucleic Acids Research. 2020;48(D1):D58-64.
[28] Dekker J, Belmont AS, Guttman M, Leshyk VO, Lis JT, Lomvardas S, et al. The 4D nucleome project. Nature. 2017;549(7671):219.
[29] Durand NC, Shamim MS, Machol I, Rao SS, Huntley MH, Lander ES, et al. Juicer provides a one-click system for analyzing loop-resolution Hi-C experiments. Cell Systems. 2016;3(1):95-8.
[30] Krietenstein N, Abraham S, Venev SV, Abdennur N, Gibcus J, Hsieh THS, et al. Ultrastructural details of mammalian chromosome architecture. Molecular Cell. 2020;78(3):554-65.
[31] Chollet F. keras. GitHub; 2015. https://github.com/fchollet/keras.
[32] Abadi M, Agarwal A, Barham P, Brevdo E, Chen Z, Citro C, et al.. TensorFlow: Large-Scale Machine Learning on Heterogeneous Systems; 2015. Available from: http://tensorflow.org/.
[33] Sima J, Chakraborty A, Dileep V, Michalski M, Klein KN, Holcomb NP, et al. Identifying cis elements for spatiotemporal control of mammalian DNA replication. Cell. 2019;176(4):816-30.
[34] McLean CY, Bristor D, Hiller M, Clarke SL, Schaar BT, Lowe CB, et al. GREAT improves functional interpretation of cis-regulatory regions. Nature Biotechnology. 2010;28(5):495-501.

### B Supplementary Figures

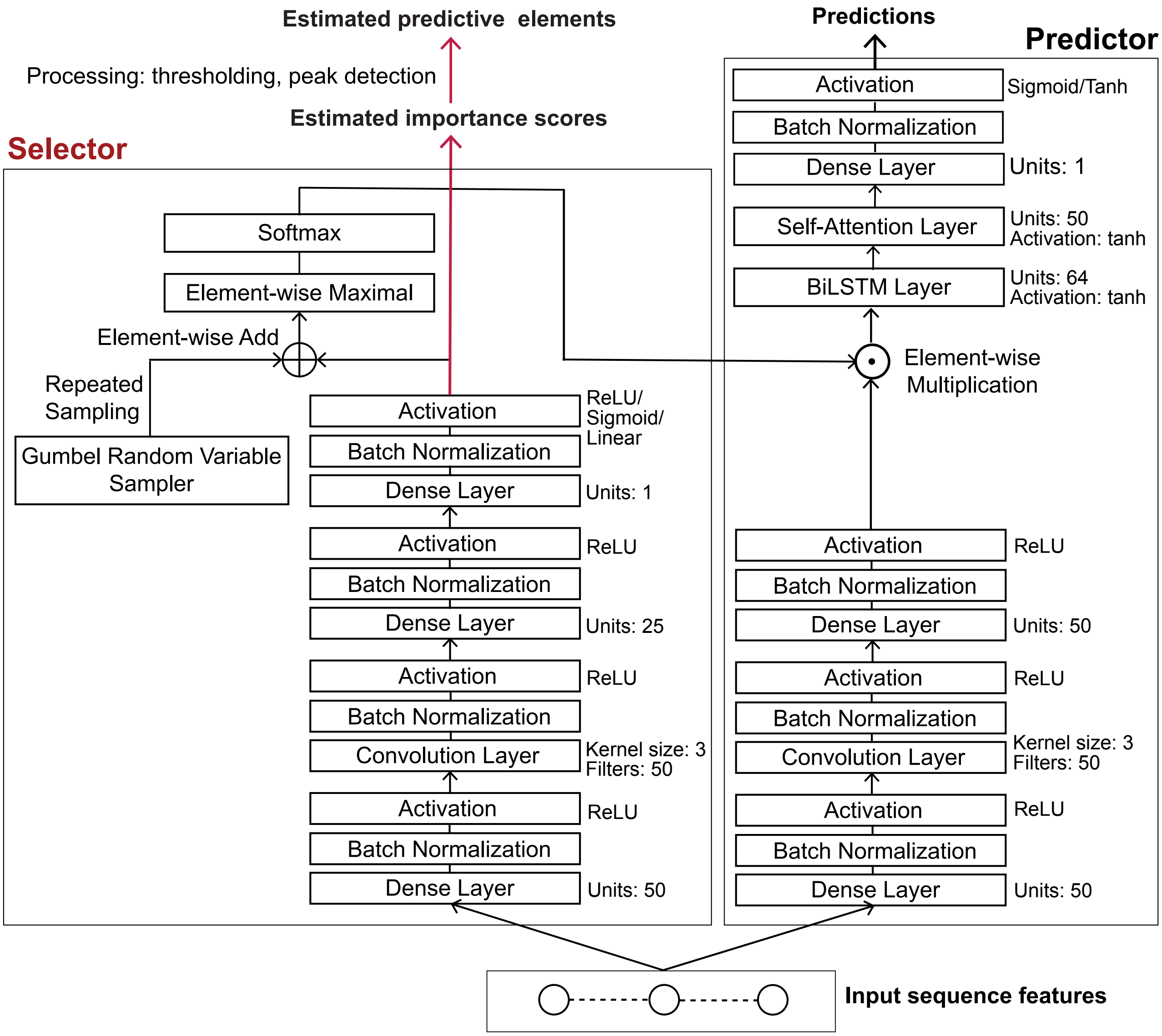


**Figure S1:** The architecture of the CONCERT model. The two primary modules are the selector and the predictor. The model is implemented using Keras/Tensorflow [31, 32]. The hyperparameter configurations used for each layer or component in practice are shown next to the corresponding layer or component.

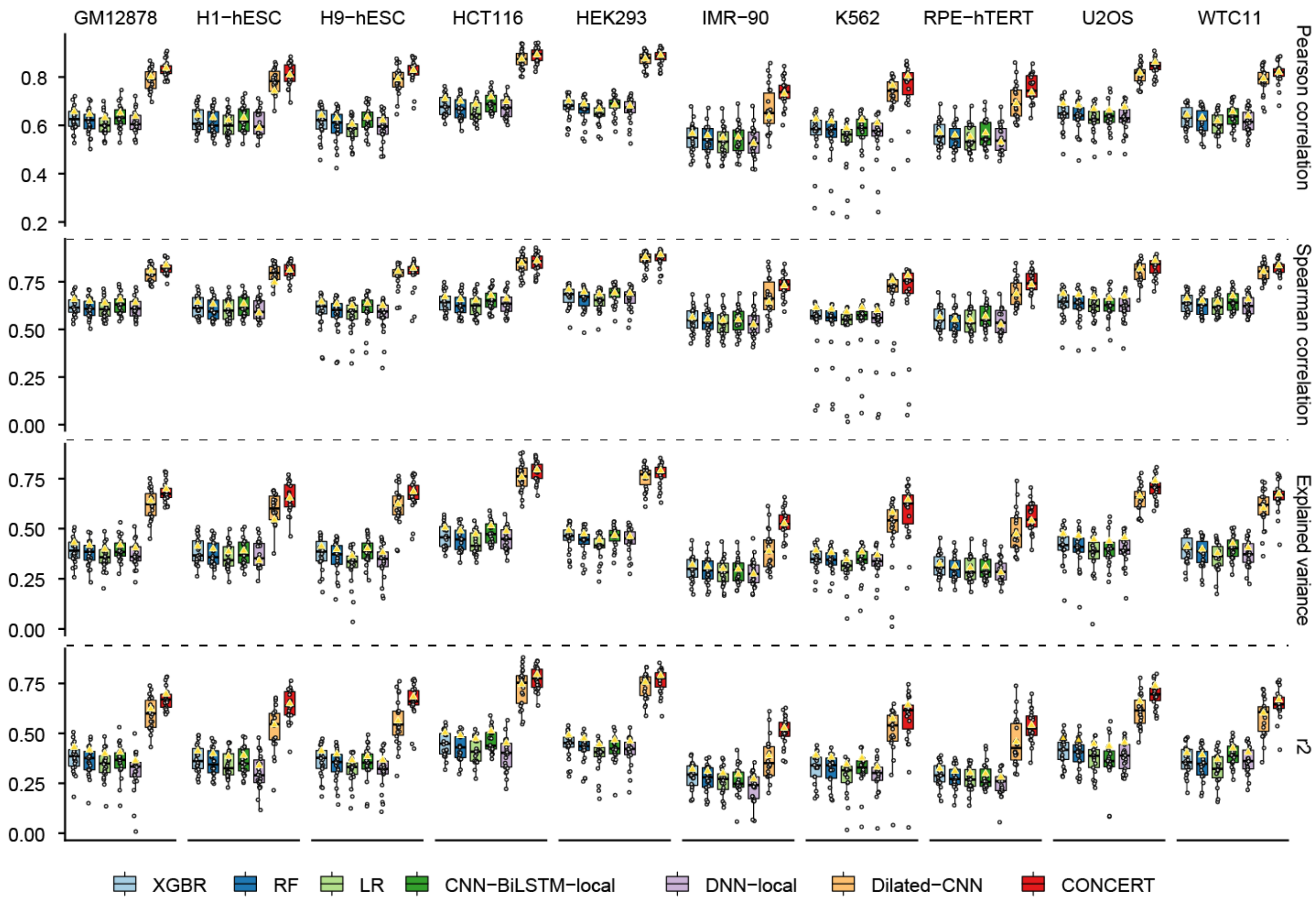


**Figure S2:** RT prediction performance measured by Pearson correlation coefficient (PCC), Spearman's rank correlation coefficient (Spearman's $\rho$), explained variance, and $R^2$ score in 10 human cell lines.

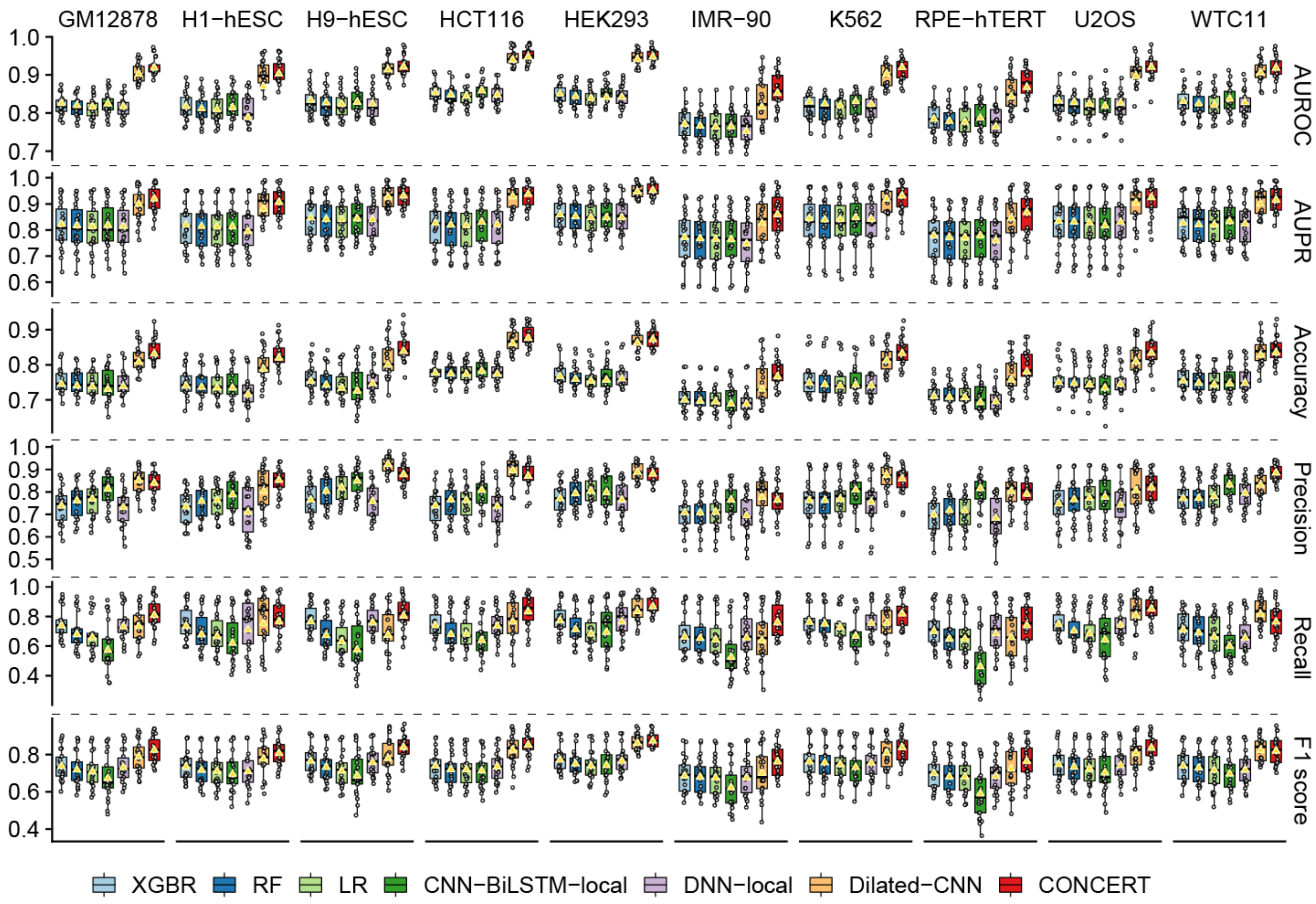


**Figure S3:** RT classification performance measured by AUROC, AUPR, accuracy, precision, recall, and $F_1$ score in 10 human cell lines.

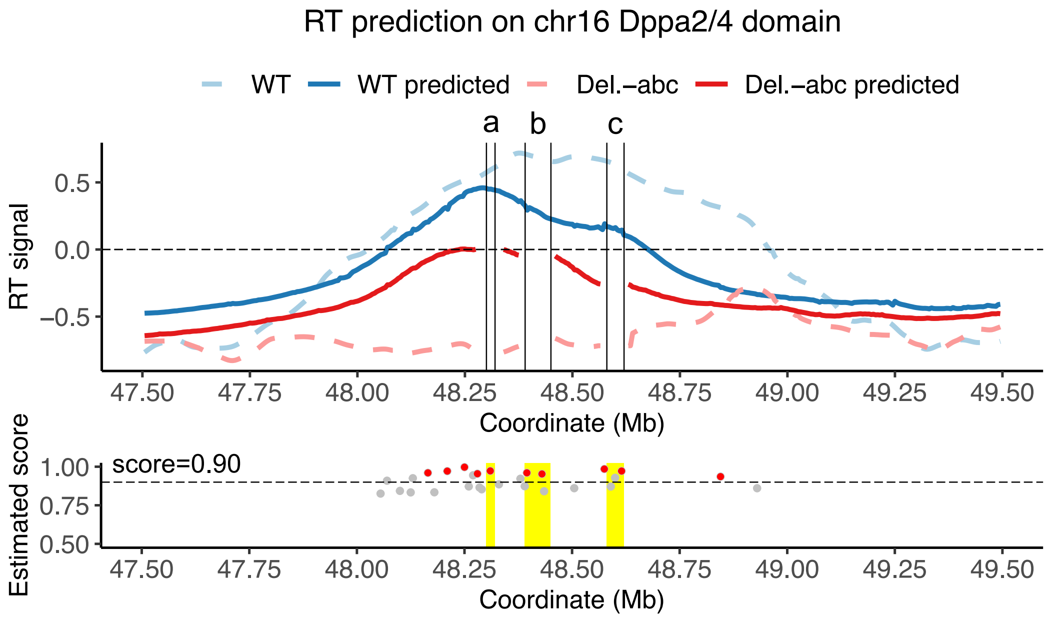


**Figure S4:** RT prediction on the *Dppa2/4* locus on chromosome 16 in mESCs [33]. The upper panel shows the CONCERT predicted signals on WT allele (without deletions) and the mutant allele (with ERCE a,b,c deleted). The bottom panel shows the normalized estimated importance scores and identified genomic loci from CONCERT. Normalized estimated scores below 0.50 are filtered. CONCERT identified important genomic loci are marked with red color. The yellow rectangles show the positions of ERCE a,b,c.

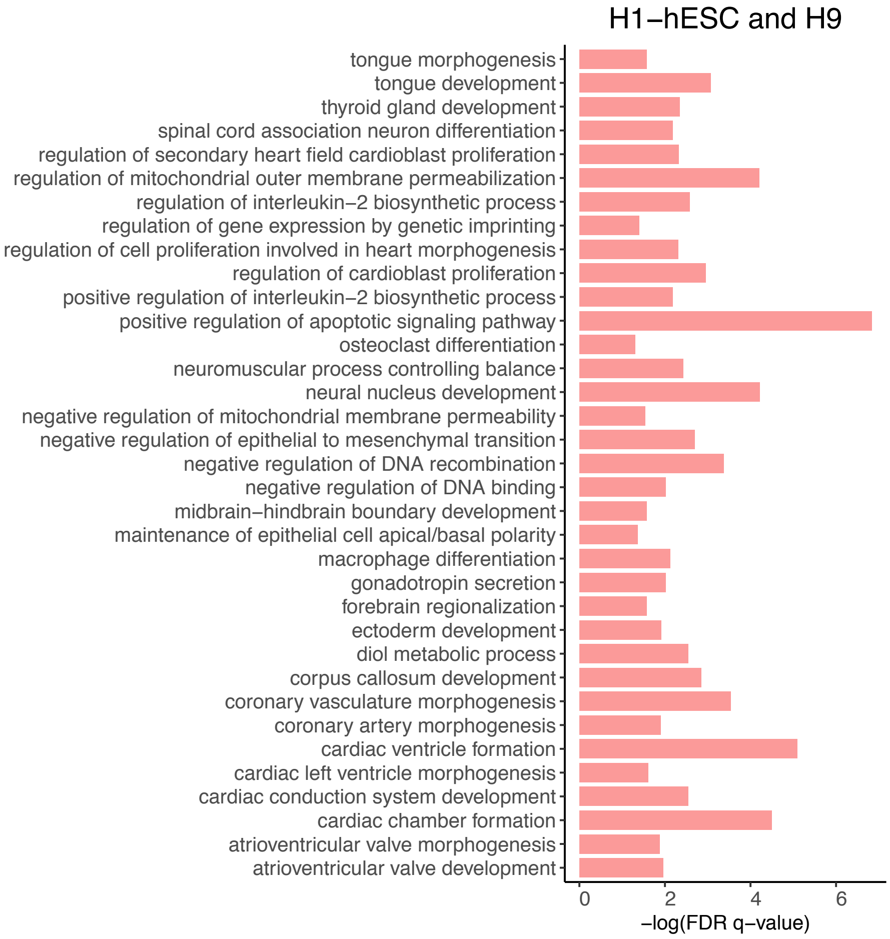


**Figure S5:** Gene Ontology (GO) analysis of the identified hESC-specific predictive genomic loci using GREAT [34]. The hESC cell types here include H1-hESC and H9-hESC. The vertical axis shows the list of gene functions that are found to be significantly associated with the identified hESC-specific predictive genomic loci. The horizontal axis shows the logarithm with base 10 (log10) value of the false positive rate (FDR, or $q$-value) based on the binomial test performed by GREAT. The FDRs of the identified associated gene functions are smaller than 0.05.

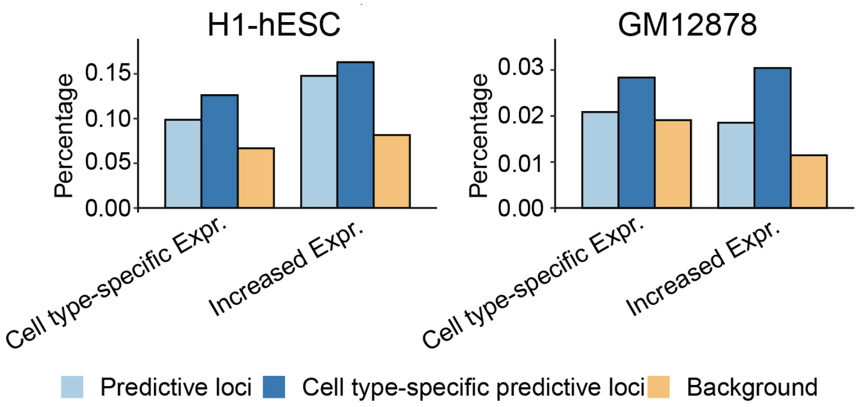


**Figure S6:** Percentage of the CONCERT predicted important loci in proximity to the genes with cell type-specific expressions or cell type-specific increased expressions in H1-hESC and GM12878.

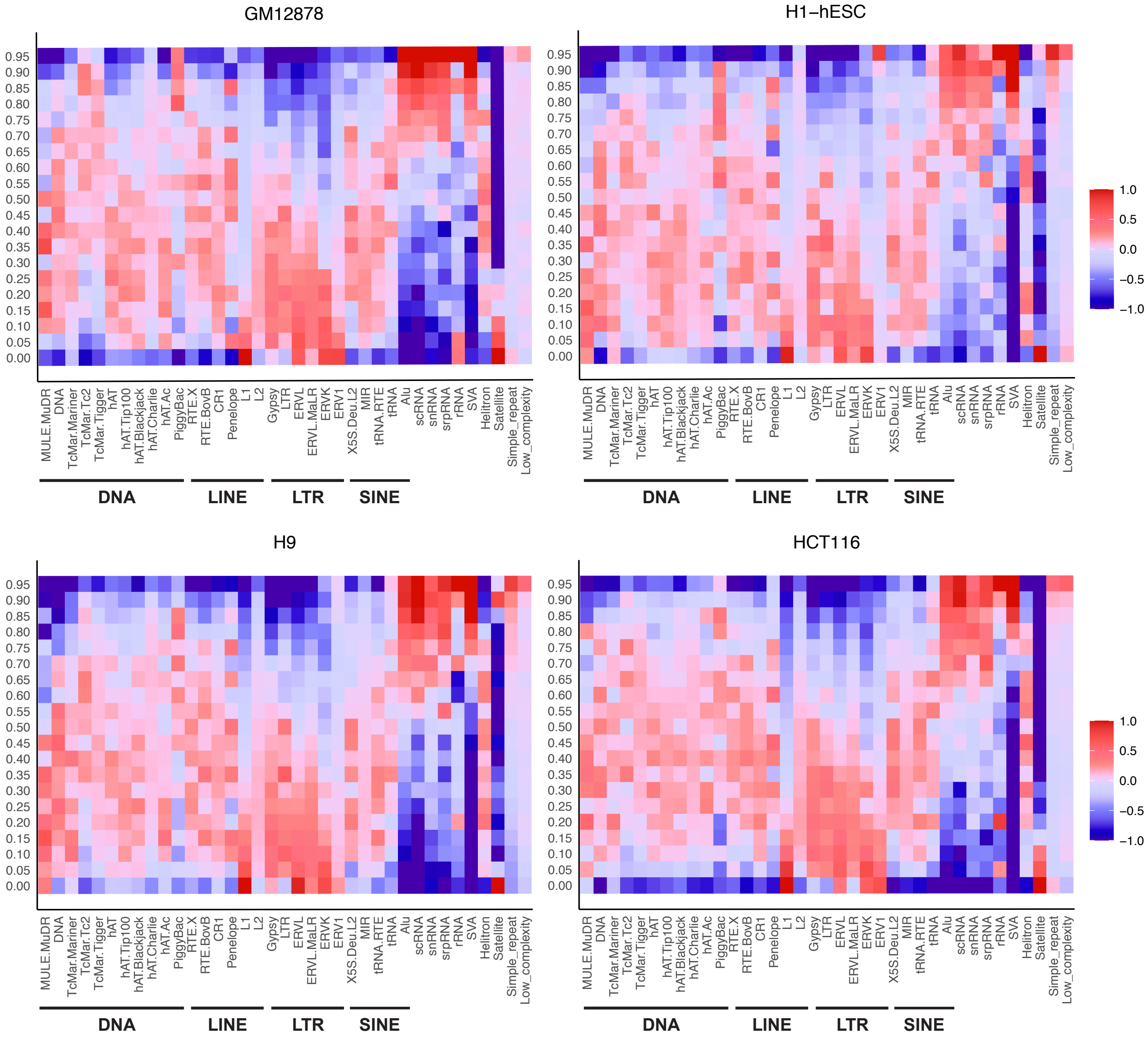


**Figure S7:** TE/RE enrichment fold change of genomic loci at different estimated sequence importance levels genome-wide in GM12878, H1-hESC, H9-hESC, and HCT116. For each cell line, we classified the genomic loci into $K = 20$ groups based on importance score ranking in descending order. Each row corresponds to a group, representing an importance score level. The first row represents genomic loci with top 5% estimated importance. Each entry represents the fold change of enrichment for the corresponding TE/RE family in each importance group in comparison with the average coverage across all groups.

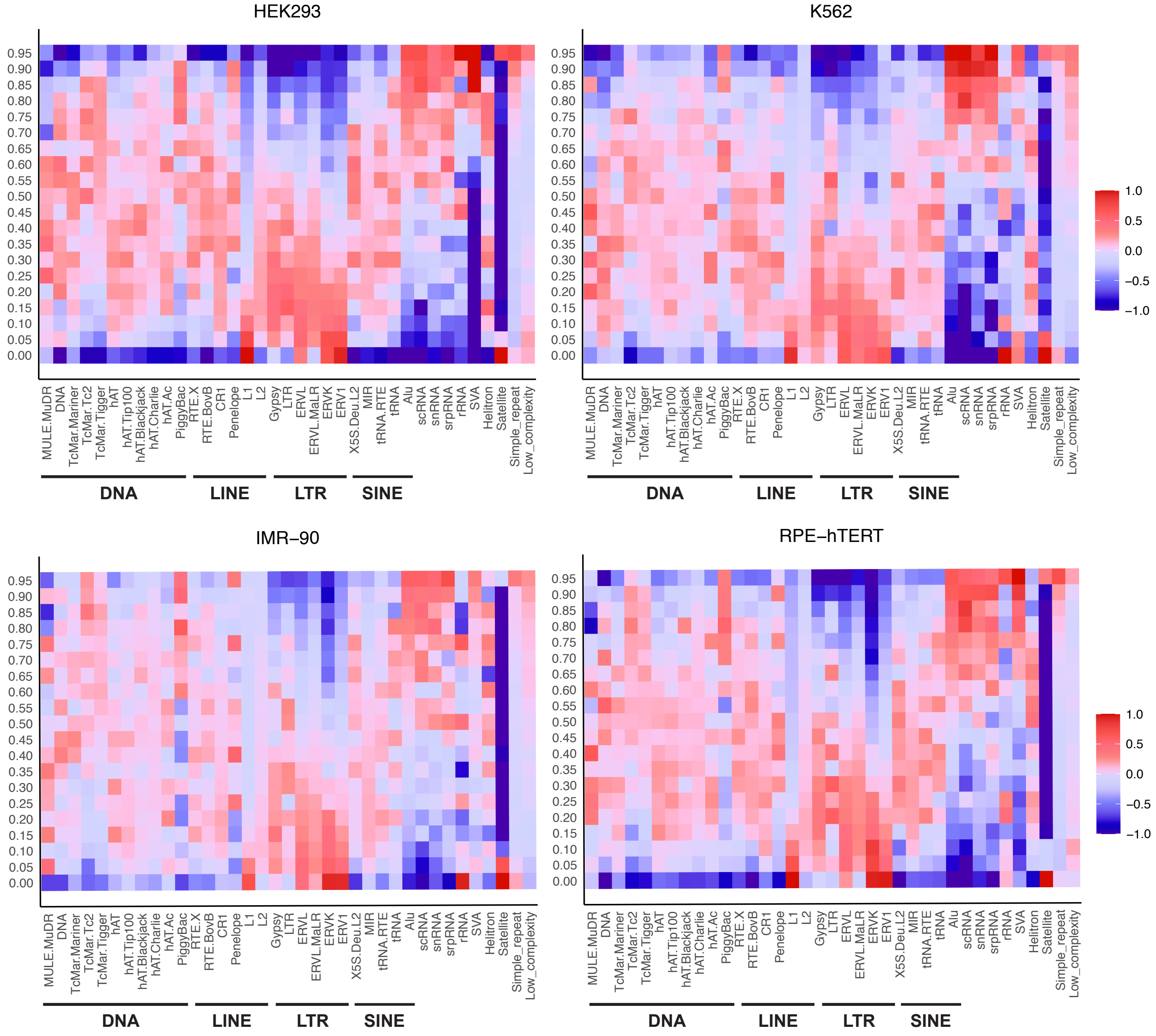


**Figure S8:** TE/RE enrichment fold change of genomic loci at different estimated sequence importance levels genome-wide in HEK293, K562, IMR-90, and RPE-hTERT. For each cell type, we classified the genomic loci into $K = 20$ groups based on importance score ranking in descending order. Each row corresponds to a group, representing an importance score level. The first row represents genomic loci with top 5% estimated importance. Each entry represents the fold change of enrichment for the corresponding RE family in each importance group in comparison with the average coverage across all groups.

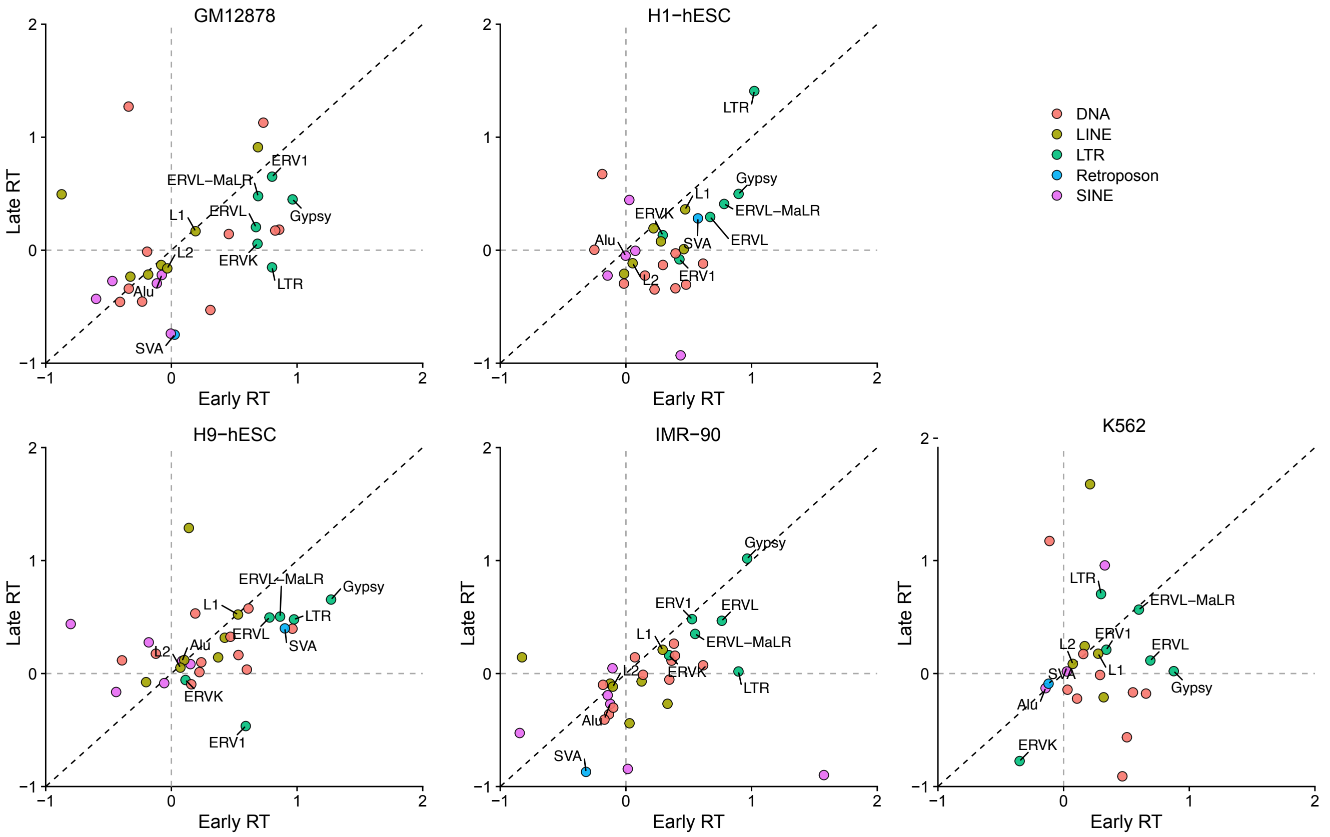


**Figure S9:** Log2 fold change of the percentage of the CONCERT predicted important genomic loci depleted of known regulatory elements and overlapping with a specific TE family vs. the percentage of predicted important loci overlapping with both known regulatory elements and the specific TE family in early and late RT regions in GM12878, H1-hESC, H9-hESC, IMR-90, and K562.

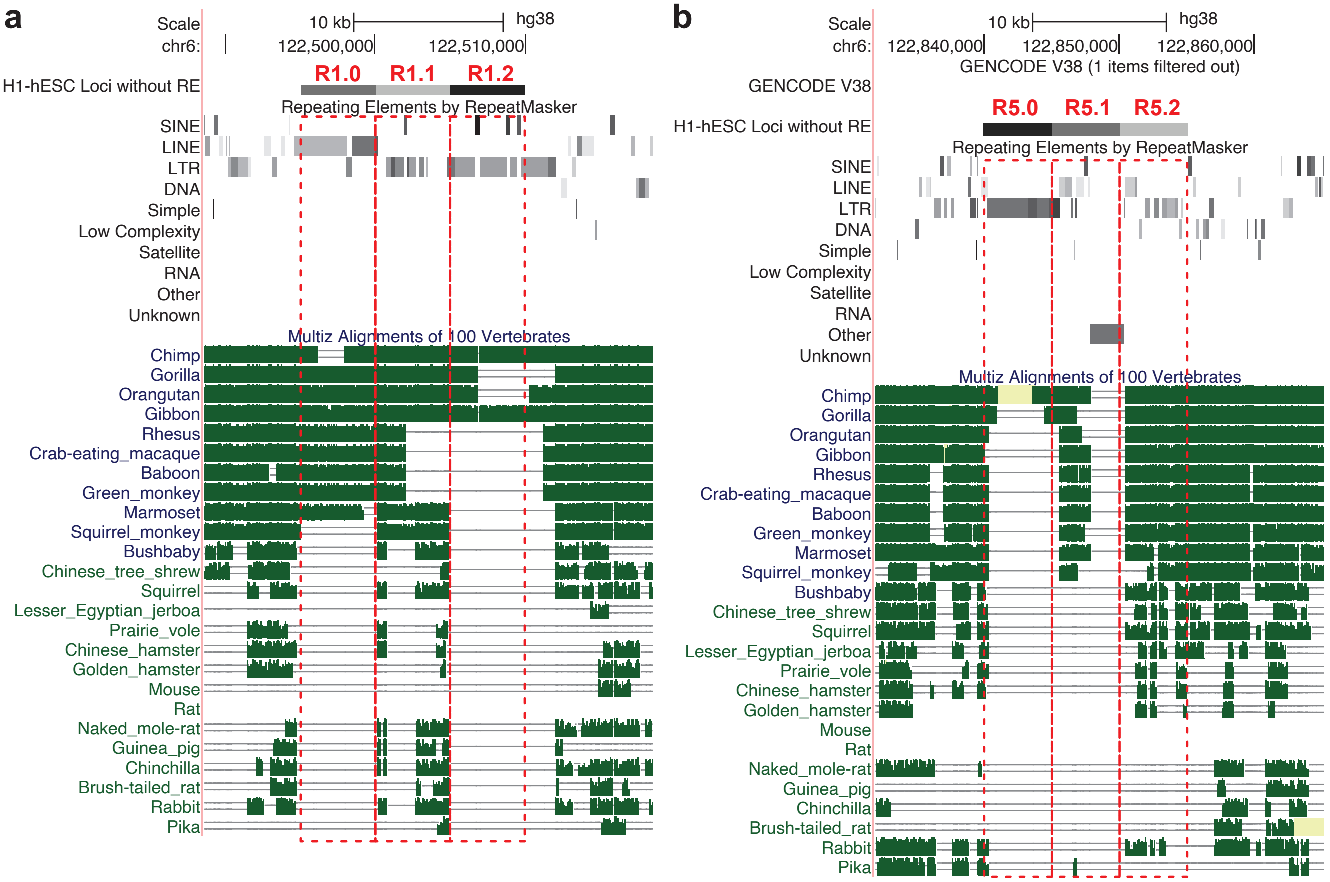


**Figure S10:** Multi-species sequence alignment at two CONCERT identified important genomic loci that are depleted with known regulatory elements and enriched with specific TEs. The loci with relatively strongest effects on RT prediction are R1.2 and R5.0.

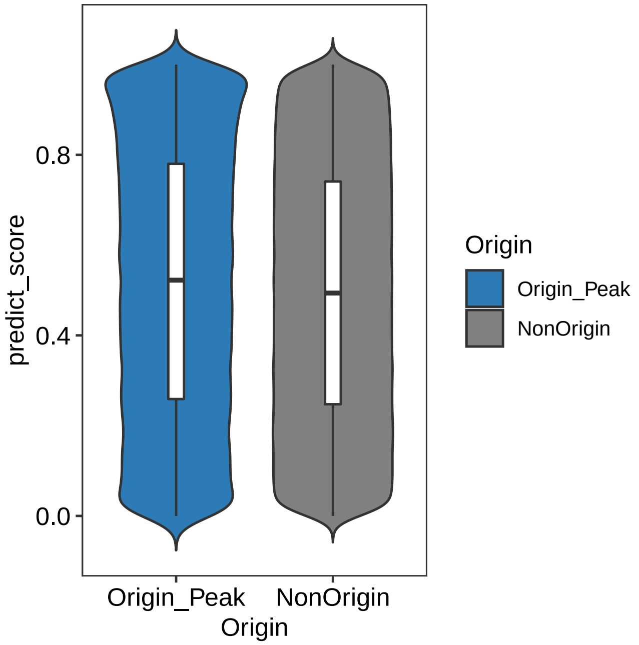


**Figure S11:** Distribution of the CONCERT estimated importance scores on RT origins and non-origins in H1-hESC. RT origins were defined by the SNS-seq peaks in H1-hESC.

### C Supplementary Tables

| Cell type | Method | PCC | Spearman's $\rho$ | Explained variance | $R^2$ score |
|---|---|---|---|---|---|
| GM12878 | XGBR | 0.6556 | 0.6573 | 0.4298 | 0.4297 |
| | RF | 0.6460 | 0.6471 | 0.4167 | 0.4166 |
| | LR | 0.6311 | 0.6387 | 0.3983 | 0.3982 |
| | DNN-local | 0.6362 | 0.6404 | 0.4039 | 0.3629 |
| | CNN-BiLSTM-local | 0.6511 | 0.6563 | 0.4178 | 0.4052 |
| | Dilated-CNN | 0.8007 | 0.8072 | 0.6399 | 0.6218 |
| | CONCERT | **0.8363** | **0.8380** | **0.6982** | **0.6968** |
| H1-hESC | XGBR | 0.6422 | 0.6427 | 0.4123 | 0.4115 |
| | RF | 0.6319 | 0.6318 | 0.3990 | 0.3984 |
| | LR | 0.6211 | 0.6268 | 0.3856 | 0.3846 |
| | DNN-local | 0.5890 | 0.5863 | 0.3398 | 0.3382 |
| | CNN-BiLSTM-local | 0.6312 | 0.6341 | 0.3896 | 0.3896 |
| | Dilated-CNN | 0.7420 | 0.7465 | 0.5444 | 0.5425 |
| | CONCERT | **0.8086** | **0.8107** | **0.6537** | **0.6481** |
| H9-hESC | XGBR | 0.6418 | 0.6449 | 0.4119 | 0.4114 |
| | RF | 0.6299 | 0.6326 | 0.3966 | 0.3963 |
| | LR | 0.6047 | 0.6232 | 0.3651 | 0.3645 |
| | DNN-local | 0.6205 | 0.6247 | 0.3813 | 0.3662 |
| | CNN-BiLSTM-local | 0.6329 | 0.6368 | 0.4003 | 0.3838 |
| | Dilated-CNN | 0.7928 | 0.8041 | 0.6226 | 0.5618 |
| | CONCERT | **0.8168** | **0.8123** | **0.6671** | **0.6611** |
| WTC11 | XGBR | 0.6430 | 0.6617 | 0.4109 | 0.4027 |
| | RF | 0.6321 | 0.6502 | 0.3985 | 0.3916 |
| | LR | 0.6177 | 0.6441 | 0.3779 | 0.3692 |
| | DNN-local | 0.6391 | 0.6572 | 0.4053 | 0.4047 |
| | CNN-BiLSTM-local | 0.6556 | 0.6753 | 0.4266 | 0.4257 |
| | Dilated-CNN | 0.7926 | 0.8043 | 0.6023 | 0.6016 |
| | CONCERT | **0.8201** | **0.8333** | **0.6726** | **0.6678** |
| HCT116 | XGBR | 0.7101 | 0.6699 | 0.5042 | 0.5040 |
| | RF | 0.6996 | 0.6575 | 0.4892 | 0.4889 |
| | LR | 0.6879 | 0.6515 | 0.4731 | 0.4728 |
| | DNN-local | 0.6984 | 0.6600 | 0.4875 | 0.4512 |
| | CNN-BiLSTM-local | 0.7176 | 0.6777 | 0.5145 | 0.5115 |
| | Dilated-CNN | 0.8742 | 0.8475 | 0.7575 | 0.7446 |
| | CONCERT | **0.8913** | **0.8598** | **0.7936** | **0.7934** |

**Table S1:** Performance evaluation of RT prediction using different methods in 10 human cell lines (part I). Performance evaluation is based on Pearson correlation coefficient (PCC), Spearman's rank correlation coefficient (Spearmans' $\rho$), explained variance and $R^2$ score between the predicted RT signals and the real RT signals across 22 autosomes in 10 human cell types using different methods for prediction. The highest performance of the compared methods is in bold font.

| Cell type | Method | PCC | Spearman's $\rho$ | Explained variance | $R^2$ score |
|---|---|---|---|---|---|
| HEK293 | XGBR | 0.7011 | 0.7083 | 0.4912 | 0.4912 |
| | RF | 0.6885 | 0.6956 | 0.4741 | 0.4740 |
| | LR | 0.6667 | 0.6851 | 0.4433 | 0.4433 |
| | DNN-local | 0.6802 | 0.6890 | 0.4617 | 0.4601 |
| | CNN-BiLSTM-local | 0.6845 | 0.6917 | 0.4663 | 0.4663 |
| | Dilated-CNN | 0.8747 | 0.8788 | 0.7606 | 0.7558 |
| | CONCERT | **0.8896** | **0.8918** | **0.7892** | **0.7889** |
| K562 | XGBR | 0.6251 | 0.6137 | 0.3908 | 0.3903 |
| | RF | 0.6172 | 0.6060 | 0.3807 | 0.3802 |
| | LR | 0.5882 | 0.5927 | 0.3455 | 0.3448 |
| | DNN-local | 0.6095 | 0.5989 | 0.3685 | 0.3307 |
| | CNN-BiLSTM-local | 0.6221 | 0.6136 | 0.3843 | 0.3762 |
| | Dilated-CNN | 0.7644 | 0.7562 | 0.5811 | 0.5729 |
| | CONCERT | **0.8032** | **0.7829** | **0.6449** | **0.6417** |
| IMR90 | XGBR | 0.5664 | 0.5631 | 0.3206 | 0.3200 |
| | RF | 0.5576 | 0.5539 | 0.3109 | 0.3102 |
| | LR | 0.5488 | 0.5481 | 0.3010 | 0.3003 |
| | DNN-local | 0.5253 | 0.5246 | 0.2739 | 0.2715 |
| | CNN-BiLSTM-local | 0.5515 | 0.5504 | 0.2982 | 0.2938 |
| | Dilated-CNN | 0.6360 | 0.6438 | 0.3884 | 0.3877 |
| | CONCERT | **0.7301** | **0.7293** | **0.5261** | **0.5261** |
| RPE-hTERT | XGBR | 0.5704 | 0.5643 | 0.3251 | 0.3223 |
| | RF | 0.5596 | 0.5527 | 0.3131 | 0.3105 |
| | LR | 0.5525 | 0.5496 | 0.3049 | 0.3020 |
| | DNN-local | 0.5302 | 0.5247 | 0.2809 | 0.2802 |
| | CNN-BiLSTM-local | 0.5679 | 0.5681 | 0.3118 | 0.2984 |
| | Dilated-CNN | 0.6902 | 0.6869 | 0.4736 | 0.4566 |
| | CONCERT | **0.7352** | **0.7334** | **0.5400** | **0.5399** |
| U2OS | XGBR | 0.6893 | 0.6880 | 0.4751 | 0.4751 |
| | RF | 0.6823 | 0.6807 | 0.4652 | 0.4652 |
| | LR | 0.6682 | 0.6712 | 0.4464 | 0.4464 |
| | DNN-Local | 0.6754 | 0.6740 | 0.4558 | 0.4545 |
| | CNN-BiLSTM-local | 0.6622 | 0.6634 | 0.4369 | 0.4356 |
| | Dilated-CNN | 0.8165 | 0.8183 | 0.6629 | 0.6597 |
| | CONCERT | **0.8607** | **0.8608** | **0.7387** | **0.7378** |

**Table S2:** Performance evaluation of RT prediction using different methods in 10 human cell lines (part II). Performance evaluation is based on Pearson correlation coefficient (PCC), Spearman's rank correlation coefficient (Spearmans' $\rho$), explained variance and $R^2$ score between the predicted RT signals and the real RT signals across 22 autosomes in 10 human cell types using different methods for prediction. The highest performance of the compared methods is in bold font.

| Chromosome | PCC | Spearman's $\rho$ | Chromosome | PCC | Spearman's $\rho$ |
|---|---|---|---|---|---|
| chr1 | 0.8505 | 0.8522 | chr12 | 0.8254 | 0.8272 |
| chr2 | 0.7719 | 0.7721 | chr13 | 0.7884 | 0.7989 |
| chr3 | 0.8016 | 0.8028 | chr14 | 0.8469 | 0.8439 |
| chr4 | 0.7903 | 0.7838 | chr15 | 0.8282 | 0.8249 |
| chr5 | 0.7701 | 0.7738 | chr16 | 0.8647 | 0.8707 |
| chr6 | 0.8063 | 0.8054 | chr17 | 0.8840 | 0.8482 |
| chr7 | 0.6940 | 0.7041 | chr18 | 0.8100 | 0.8053 |
| chr8 | 0.7462 | 0.7300 | chr19 | 0.7484 | 0.8005 |
| chr9 | 0.8110 | 0.8153 | chr20 | 0.7986 | 0.8123 |
| chr10 | 0.7794 | 0.7808 | chr21 | 0.8664 | 0.8582 |
| chr11 | 0.8657 | 0.8696 | chr22 | 0.8577 | 0.7671 |

**Table S3:** RT prediction performance of CONCERT on each autosomal chromosome in H1-hESC. Performance evaluation is based on Pearson correlation coefficients (PCC) and Spearman's rank correlation coefficient (Spearman's $\rho$) between the predicted RT signals and the real RT signals on each autosome in H1-hESC.

| Cell type | RT Fraction | Accession Number |
|---|---|---|
| H1-hESC | Early | 4DNES4B7RLAV |
| H1-hESC | Late | 4DNESUJC9Y83 |
| H9-hESC | Early | 4DNESJQH3HJ9 |
| H9-hESC | Late | 4DNEST651P8O |
| GM12878 | Early | 4DNESO83H9ZI |
| GM12878 | Late | 4DNESDQ9PZOX |
| HCT116 | Early | 4DNESSYKYIK3 |
| HCT116 | Late | 4DNESLC8TDK4 |
| HEK293 | Early | 4DNESH4XLJCW |
| HEK293 | Late | 4DNESSV33VOL |
| K562 | Early | 4DNES9YA22WT |
| K562 | Late | 4DNES1GSPUT8 |
| IMR-90 | Early/Late | ENCSR734AQN |
| RPE-hTERT | Early | 4DNESUNOW1OZ |
| RPE-hTERT | Late | 4DNES674QWXK |
| U2OS | Early | 4DNES1P18J2X |
| U2OS | Late | 4DNES99LXRYK |
| WTC11 | Early | 4DNESGWKYYWO |
| WTC11 | Late | 4DNES4WLR8MV |

**Table S4:** E/L 2-fraction Repli-Seq datasets used in this paper were downloaded from 4DN Data Portal. The Repli-seq for IMR-90 was downloaded from the ENCODE Project Data Portal.

| Input feature type | PCC | Spearman's $\rho$ | Explained variance | $R^2$ score |
|---|---|---|---|---|
| Feature I | 0.8086 | 0.8107 | 0.6537 | 0.6481 |
| Feature II | 0.7827 | 0.7811 | 0.5970 | 0.5652 |
| GC-based feature | 0.7250 | 0.7190 | 0.5017 | 0.4369 |
| Feature I+phyloP score | 0.8030 | 0.8012 | 0.6495 | 0.6487 |

**Table S5:** Performance evaluation based on Pearson correlation coefficient (PCC), Spearman's rank correlation coefficient (Spearmans' $\rho$), explained variance and $R^2$ score between the predicted RT signals and the real RT signals across 22 autosomes in H1-hESC using different types of features with the basic CONCERT model. Feature I: combination of *K*-mer frequency features, GC-based features, Feature II: combination of *K*-mer frequency features, GC-based features, and TF binding motif features, Feature I+ phyloP score: combination of *K*-mer frequency features, GC-based features, and phyloP score features.

| Model type | PCC | Spearman's $\rho$ | Explained variance | $R^2$ score |
|---|---|---|---|---|
| CONCERT-basic | 0.8086 | 0.8107 | 0.6537 | 0.6481 |
| CONCERT-hierarchical | 0.7850 | 0.7911 | 0.6220 | 0.6336 |

**Table S6:** Performance evaluation based on Pearson correlation coefficient (PCC), Spearman's rank correlation coefficient (Spearman's $\rho$), explained variance and $R^2$ score between the predicted RT signals and the real RT signals across 22 autosomes in H1-hESC using the basic CONCERT model with predefined feature representation Feature I (combination of *K*-mer frequency features and GC-based features) and the CONCERT-hierarchical model.

| Cell type | GM12878 | H1-hESC | H9-hESC | HCT116 | HEK293 |
|---|---|---|---|---|---|
| Number of identified predictive elements | 5037 | 5466 | 5477 | 5001 | 5459 |
| Cell type | K562 | IMR-90 | RPE-hTERT | U2OS | |
| Number of identified predictive elements | 5453 | 5108 | 5460 | 3059 | |

**Table S7:** The number of identified predictive elements in each cell type that are also identified in at least another four cell types with flanking region size of +/-2 bins (bin size: 5Kb). An identified predictive element is an identified predictive genomic locus or a segment merging identified predictive genomic loci that are less than 5 bins apart. If two identified predictive elements/loci with flanking regions in two cell types overlap with each other, we consider each of the the elements as also identified in the other cell type.

| Cell type | GM12878 | H1-hESC | H9-hESC | HCT116 | HEK293 |
|---|---|---|---|---|---|
| Number of identified predictive elements | 9041 | 9659 | 8958 | 7307 | 9098 |
| Cell type | K562 | IMR-90 | RPE-hTERT | U2OS | |
| Number of identified predictive elements | 12779 | 14495 | 10145 | 11982 | |

**Table S8:** The number of predictive elements that are only identified in one of the nine human cell types with flanking region size of +/-2 bins.
